# Estimation of site frequency spectra from low-coverage sequencing data using stochastic EM reduces overfitting, runtime, and memory usage

**DOI:** 10.1101/2022.05.24.493190

**Authors:** Malthe Sebro Rasmussen, Genís Garcia-Erill, Thorfinn Sand Korneliussen, Carsten Wiuf, Anders Albrechtsen

## Abstract

The site frequency spectrum (SFS) is an important summary statistic in population genetics used for inference on demographic history and selection. However, estimation of the SFS from called genotypes introduce bias when working with low-coverage sequencing data. Methods exist for addressing this issue, but sometimes suffer from two problems. First, they can have very high computational demands, to the point that it may not be possible to run estimation for genome-scale data. Second, existing methods are prone to overfitting, especially for multi-dimensional SFS estimation. In this article, we present a stochastic expectation-maximisation algorithm for inferring the SFS from NGS data that addresses these challenges. We show that this algorithm greatly reduces runtime and enables estimation with constant, trivial RAM usage. Further, the algorithm reduces overfitting and thereby improves downstream inference. An implementation is available at github.com/malthesr/winsfs.

## 1 Introduction

The site frequency spectrum (SFS) is the joint distribution of allele frequencies among one or more populations, and it serves as an important summary statistic in population genetics. For instance, the SFS is sufficient for computing nucleotide diversity [1], F_st_ [2], and *f*-statistics [3]. Furthermore, the SFS may be used for inferring demographic history [4–6] and selection [7–9].

When working with high-quality data, it is usually straightforward to estimate the SFS from called genotypes. However, when genotype calls are uncertain, standard methods lead to significant bias in the estimated SFS [10], which propagates to downstream inference [11]. In particular, this situation arises when working with next-generation sequencing (NGS) data at low coverage and may be compounded by additional data-quality issues. Low-coverage NGS data is sometimes the only available option, for instance when working with ancient DNA [12–14]. Sequencing at low coverage is also a popular choice to reduce sequencing costs, since most of the key population genetics analysis remain possible with such data [15].

To estimate the SFS from low-coverage data, several methods have been proposed which account for the genotype uncertainty in estimation of the SFS [10, 16]. These are based on finding the SFS that maximises the data likelihood using numeric optimisation. Two factors combine to create a computational challenge for such methods. First, in order to achieve an accurate estimate of the SFS, these methods usually require many iterations, each of which requires a full pass over the input data. Second, unlike most genetics analyses, the SFS cannot be based on only the small subset of the variable sites, but must consider all sites. Taken together, this means that some summary of the full data must be held in RAM and iterated over many times. For genome-scale NGS data from more than a few dozen samples, or in more than one dimension, this is often not computationally feasible, as tens of hours of runtime and hundreds of gigabytes of RAM may be required. Current approaches for dealing with this issue restrict the analysis to fewer individuals and/or smaller regions of the genome [17], leading to less accurate results.

An additional problem with current methods is that they are prone to overfitting. In the multi-dimensional setting in particular, there is often very little information available for many of the entries in the frequency spectrum. Therefore, by considering the full data set, existing algorithms risk fitting noise, leading to estimates with poor generalisability.

In this paper, we present a novel version of the stochastic expectation-maximisation (EM) algorithm for estimation of the SFS from NGS data. In each pass through the data, this algorithm updates the SFS estimate multiple times in smaller blocks of sites. We show that for low-coverage whole-genome sequencing (WGS) data, this algorithm requires only a few full passes over the data. This considerably decreases running time, and means that it is possible to estimate the SFS using constant, negligible RAM usage by streaming data from disk. Moreover, by only considering smaller subsets of the data at a time, we show that this method reduces overfitting, which in turns leads to improved downstream inference.

### 2 Methods

Estimation of the SFS from low-coverage sequencing data requires pre-computing site allele frequency likelihoods for each site, and these are based on genotype likelihoods. We begin by briefly reviewing these concepts.

### Genotype likelihoods

Assume we have NGS data *X* sampled from *K* different populations (indexed by *k*), with *N*_*k*_ individuals in the *k*th population. Further, say that we have *M* diallelic sites (indexed by *m*), so that *G*_*mkn*_ ∈ {0, 1, 2} is the genotype of a diploid individual *n* at site *m* in population *k*, coding genotypes as the number of derived alleles. In the same way, we use *X*_*mkn*_ to refer to the sequencing data at this location.

We define the genotype likelihood *p*(*X*_*mkn*_ | *G*_*mkn*_) as the probability of the data given a particular genotype. Genotype likelihoods form the basis of genotype calling and are calculated from aligned sequencing reads by various bioinformatic tools including `bcftools`/`samtools` [18, 19], `GATK` [20], and `ANGSD` [21], using slightly different models. For clarity, we outline the basic `GATK` model below, though the choice of model is not important for our purposes.

For *D* sequencing reads aligned to position *m* for individual *n* in population *k*, let *b*_*d*_ be the base call of the *d*th read. Assuming independence of base calls, we have

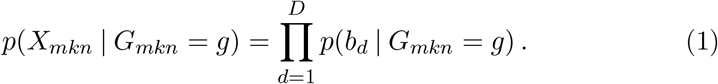

If we consider the genotype as two alleles *a*_1_, *a*_2_ ∈ {0, 1} such that *G*_*mkn*_ = *a*_1_ + *a*_2_, then by random sampling of the parental alleles,

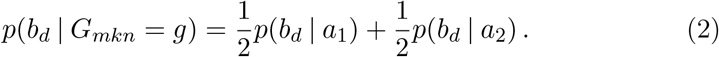

In turn, this probability is modelled by

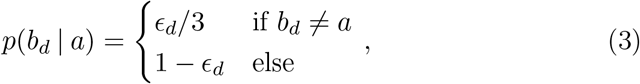

where *ϵ*_*d*_ is the sequencing error probability associated with the *d*th base.

### Site allele frequency likelihoods

Using genotype likelihoods, we can calculate site allele frequency (SAF) likelihoods, also sometimes known as sample allele frequency likelihoods. It is possible to think of the SAF likelihoods as the generalisation of genotype likelihoods from individuals to populations: instead of asking about the probability of the data for one individual given a genotype, we ask about the probability of the data for a population given the sum of their derived alleles.

More formally, define the sum of derived alleles for population *k* at site *m*,

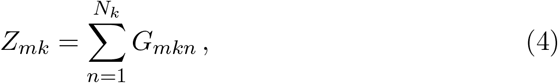

with *Z*_*mk*_ ∈ {0, 1, …, 2*N*_*k*_} each corresponding to possible sample frequencies {0, 1*/*2*N*_*k*_, …, 1}. Now define the SAF likelihood for a single population *k*,

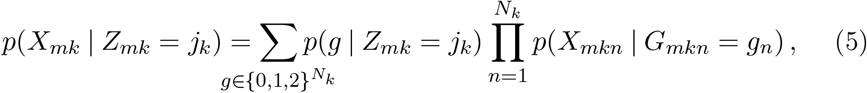

where *X*_*mk*_ is the data for all individuals sampled in population *k* at site *m, p*(*g* | *Z*_*mk*_ = *j*_*k*_) is the combinatorial probability of the genotype vector 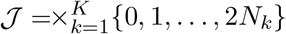 conditional on the sum of the genotypes being *j*_*k*_, and *p*(*X*_*mkn*_ | *G*_*mkn*_ = *g*_*n*_) is a standard genotype likelihood. Using a dynamic programming algorithm, SAF likelihoods can be calculated from the genotype likelihoods of *N* individuals in *O*(*N*^2^) time per site [22], and a linear time approximation has also been given [23].

To extend this to the multi-dimensional SFS with *K* populations, let 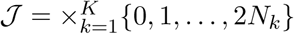 be the set of possible derived allele count combinations across populations, let *X*_*m*_ be the data across all individuals in all populations at site *m*, and define *Z*_*m*_ = (*Z*_*m*1_, …, *Z*_*mK*_) ∈ 𝒥. Then

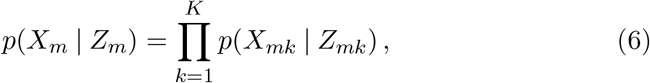

is the joint SAF likelihood for *K* populations.

### Site frequency spectrum

Using the definition of 𝒥 above, we define the SFS as a parameter *ϕ* = {*ϕ*_*j*_ : *j* ∈ 𝒥} such that *ϕ*_*j*_ is the probability that *Z*_*m*_ = *j*. That is,

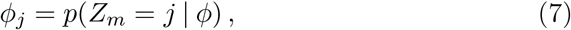

for site *m*. That is, the SFS is the probability of a particular vector of derived allele sums at a site chosen at random.

When genotypes are available, the SFS can be estimated simply by counting observed allele count combinations. When genotypes cannot be called, the standard approach is maximum-likelihood estimation.

Assuming independence of sites, we write the likelihood function

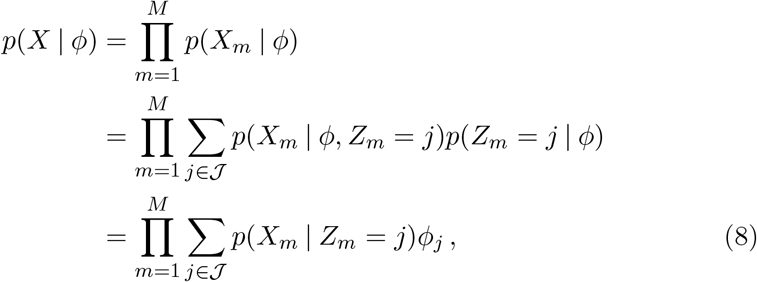

where *X*_*m*_ refers to all sequencing data for site *m*. Note that the likelihood can be expressed solely in terms of joint SAF likelihoods.

The maximum likelihood estimate 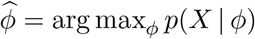 cannot be found analytically. Instead, 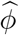 is typically estimated using some iterative procedure such as BFGS [22] or an EM algorithm [16, 21]), of which the latter has become the standard choice. An overview of the this algorithm is given below. For details and proof, see supplementary text S1.

### Standard EM algorithm

Before optimization, we pre-compute the SAF likelihoods for all sites, populations, and possible sample frequencies. In addition, we make an arbitrary initial guess of the SFS 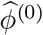. The EM algorithm then alternates between an E-step, and an M-step.

The E-step consists of computing posterior probabilities of derived allele counts conditional on the current SFS estimate,

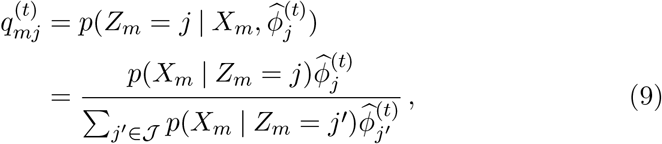

for all sites *m* ∈ {1, …, *M*} and possible derived allele counts *j* ∈ J. Note that this conditional posterior depends only on the current SFS estimate and the (joint) SAF likelihoods.

Using the result of the E-step, the M-step updates the estimate by setting

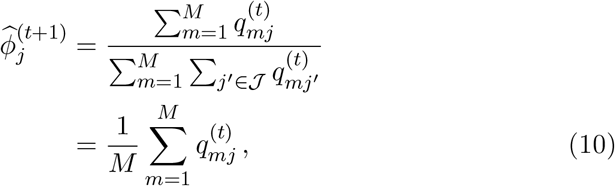

for all *j* ∈ 𝒥.

The EM algorithm guarantees a monotonically increasing likelihood of successive values number of 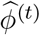. The runtime of the algorithm is linear in the iterations required before convergence, with each of iteration taking 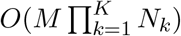 time. In practice, the standard implementation is `realSFS` [22] from the software suite `ANGSD` [21] which uses a generic EM acceleration scheme [24]. The details of this acceleration will not be important in this context, so we omit the details.

### Window EM algorithm

As in standard EM, we pre-compute all SAF likelihoods and make an arbitrary initial guess 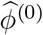 of the SFS. In addition, we choose two hyperparameters *B* (the number of blocks) and *W* (the window size). Before starting optimization, all sites indices are randomly assigned to one of *B* blocks 𝔅 = (𝔅_1_, …, 𝔅_*B*_) with |𝔅_*b*_| = ⌊*M/B*⌋ for *b < B*, and |𝔅_*B*_| = *M* mod *B*. The reason for doing so is simply to break patterns of linkage disequilibrium in particular blocks of input data, which will make the SFS within each block more similar to the global SFS. Blocks are non-overlapping and exhaustive, so that 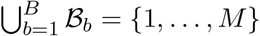 and 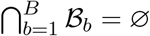.

After this initialisation, the window EM algorithm is defined as an iterative procedure that alternates between an E-step and an M-step, where the M-step in turn is split into an M1-step and an M2-step.

The E-step of the algorithm involves computing posteriors conditional on the current estimate of the SFS, much like standard EM. The difference is that we only process a single block of sites. Let *f* (*t*) = (*t* − 1) mod *B* + 1, so that *f* (1 + *xB*) = 1, *f* (2 + *xB*) = 2, … for *x* ≥ 0. Then, at time step *t*, we compute 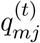 for all *m* ∈ 𝔅_*f* (*t*+1)_ and all possible derived allele counts *j* ∈ 𝒥 using eq. (9).

In the M1-step, the *q*s for the current block are used to give a block SFS estimate 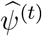. This is analogous to the standard M-step eq. (10), so that for each *j* ∈ 𝒥.

#### Algorithm 1

Window EM algorithm

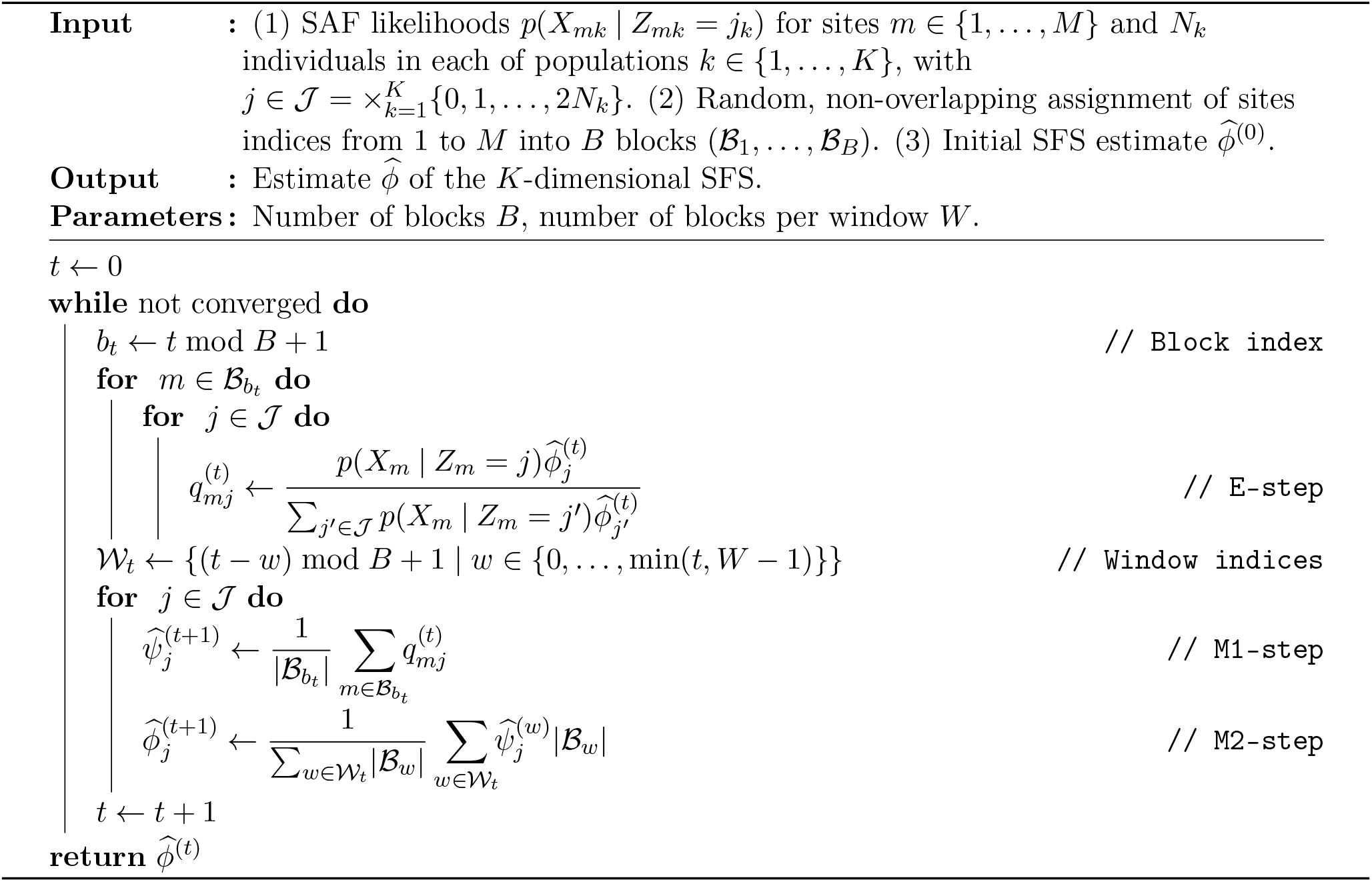

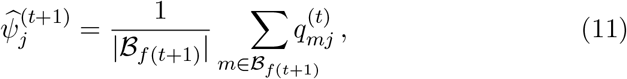

These block estimates are then used in the M2-step to update the overall SFS estimate for each *j* ∈ 𝒥,

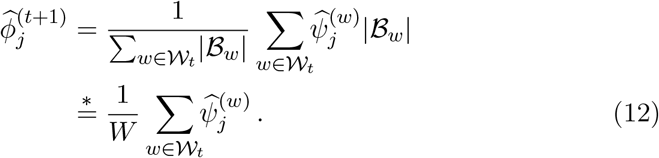

where 𝒲_*t*_ = {*f* (*t* + 1 − *w*) | *w* ∈ {0, …, min(*t, W* − 1)}} is the window of the *W* latest block indices at time *t*. We use 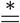 to express equality under the common special case when either *M/B* = 0 or *B* ∉ 𝒲_*t*_, so that there are no issues with blocks of unequal sizes in the current window. In this case, the M2-step simplifies to the mean of the past *W* block estimates.

Pseudo-code for window EM is given in algorithm 1, and an illustration comparing window EM to standard EM is shown in figure 1.

**Figure 1:**
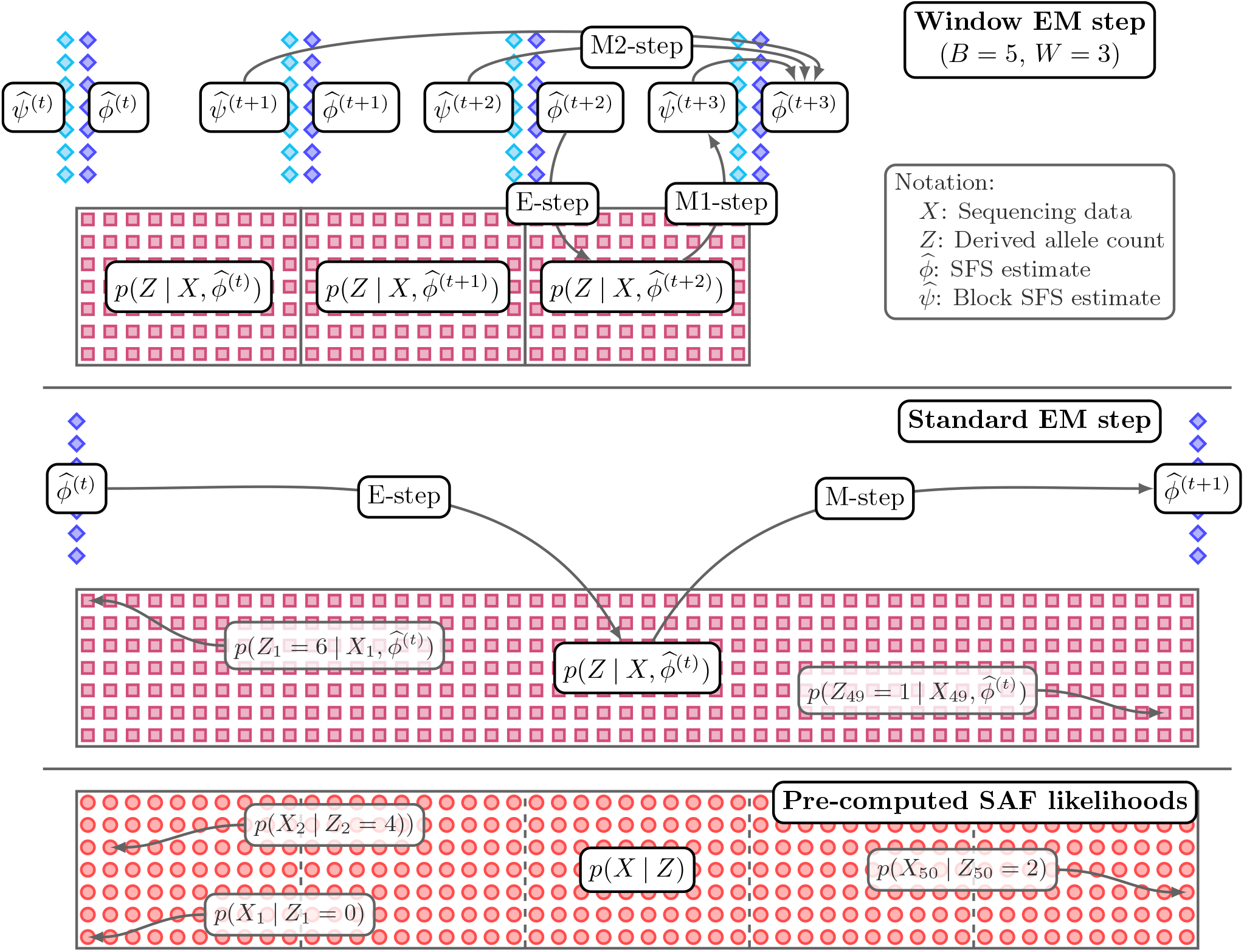
Schematic illustration of the standard and window EM algorithms for input consisting of a single population with N = 3 individuals and M = 50 sites. Sites are shown horizontally, derived allele frequencies are shown vertically. The pre-computed SAF likelihoods are illustrated at the bottom with blocks indicated by dashed lines. Standard EM computes the conditional posterior derived allele counts over all sites (E-step) and uses these to update the SFS estimate (M-step). Window EM computes the conditional posteriors for a small blocks of sites (E-step), computes a block SFS estimate after each block (M1-step), and updates the overall estimates as sliding window average (M2-step) of the W past block estimates. In this example, the sites have been split into B = 5 blocks with 10 sites each, and the sliding window covers W = 3 blocks.

In the below, we are interested in comparing standard EM and window EM. For clarity, we will use the term ‘epoch’ to refer to a full pass through the data for either algorithm. In the case of standard EM, an epoch is simply a single iteration; for window EM, an epoch corresponds to *B* iterations.

### Convergence

In the standard EM algorithm, the data log-likelihood (8) can typically be evaluated with little computational overhead during the E-step. Therefore, a common convergence criterion is based on the difference between the log-likelihood values of successive epochs. That is, let

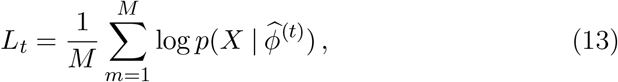

and convergence is reached when *L*_*t*+1_ − *L*_*t*_ < *δ*, for some tolerance *δ* decided ahead of time.

For window EM, the same does not apply, since no full E-step is ever taken. However, the likelihood for each block can be calculated cheaply during each block E-step. Therefore, we define for epoch *e* ∈; {1, 2, …},

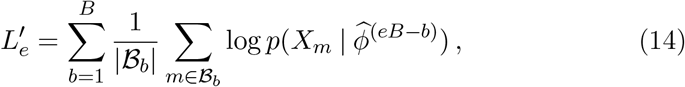

that is, the sum of log-likelihoods of SFS estimates used over the past epoch, each evaluated in the block for which they were used in a block E-step, normalised by block size for convenience. We then propose the simple convergence criterion for window EM such that convergence is defined as 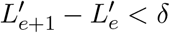.

## 3 Results

To test the window EM algorithm, we implemented it in the `winsfs` program, available at github.com/malthesr/winsfs. We compare `winsfs` to `realSFS`, which implements the standard EM algorithm and serves as the current state of art. We adopt two complementary approaches for evaluating performance of `winsfs`. First, we use two different real-world WGS data sets to compare `winsfs` to `realSFS`, which implements the standard EM algorithm and serves as the current state of the art. `realSFS` has already been validated on simulated data [21, 23], and use split training and test data sets to evaluate any observed differences. Second, we use simulated data to validate `winsfs` under conditions of known truth across a range of data qualities and sample sizes.

### Real-world data sets

We tested `winsfs` and `realSFS` on two real-world WGS data sets of very different quality as described below. An overview is shown in table 1.

**Table 1:** Overview of the input training data.

| Population | Individuals | Sites | Median depth (range) | Contigs $\geq 100$ kb | $n_{50}$ | $F_{ST}$ |
| --- | --- | --- | --- | --- | --- | --- |
| **Human** |  |  |  |  |  |  |
| YRI | 10 | $1.17 \cdot 10^9$ | $5.0\times (3.2\times\text{--}7.2\times)$ | 52 | $1.5 \cdot 10^8$ | 0.13 |
| CEU | 10 | $1.17 \cdot 10^9$ | $6.2\times (2.9\times\text{--}8.0\times)$ | | | |
| **Impala** |  |  |  |  |  |  |
| Masai Mara | 12 | $6.34 \cdot 10^8$ | $2.8\times (1.4\times\text{--}10.2\times)$ | 7811 | $3.4 \cdot 10^5$ | 0.24 |
| Shangani | 8 | $6.34 \cdot 10^8$ | $2.9\times (2.6\times\text{--}16.8\times)$ | | | |

We first analyse 10 random individuals from each of the YRI (Yoruba Nigerian) and CEU (Europeans in Utah) populations from the 1000 Genomes Project [25]. This human data was sequenced to 3 x–8 x coverage and mapped to the high quality human refence genome. We created SAF files using `ANGSD` [21] requiring minimum base and mapping quality 30 and polarising the spectrum using the chimpanzee as an outgroup. We then split this input data into test and training data, such that the first half of each autosome was assigned to the training set, and the second half to the test set. The resulting training data set contains 1.17 · 10^9^ sites for both YRI and CEU, while the test data set contains 1.35 · 10^9^ sites for both. Training set depth distributions for each individual are shown in supplementary figure 1.

We also analyse a data set of much lower quality from 12 and 8 individuals from two impala populations that we refer to as ‘Maasai Mara’ and ‘Shangani’, respectively, based on their sampling locations. These populations were sequenced to only 1 x–3 x with the addition of a single high-depth sample in each population (see supplementary figure 2). The data was mapped to a very fragmented assembly, and then we split the data into training and test sets just as for the human data. However, due to the low quality assembly we analysed only sites on contigs larger than 100 kb, and filtering sites based on depth outliers, excess heterozygosity, mappability, and repeat regions. We polarised using the impala reference itself. This process is meant to mirror a realistic workflow for working with low-quality data from a non-model organism. The impala input data ends up somewhat smaller than the human data set, with approximately 6.3 · 10^8^ sites in both test and training data sets.

Broadly, the human data is meant to exemplify medium-quality data with coverage towards the lower end, but with no other significant issues. The impala data, on the other hand, represents low-quality data: not only is the coverage low and fewer sites are available, but the impala reference genome is poor quality with 7811 contigs greater than 100 kb and *n*_50_ = 3.4 · 10^*−*5^ (that is, 50 % of the assembly bases lie on contigs of this size or greater). This serves to introduce further noise in the mapping process, which amplifies the overall data uncertainty. Finally, the impala populations are more distinct, with F_st_ ≈ 0.24 compared to 0.13 between the human populations. As we will see below, this creates additional challenges for estimation of the two-dimensional SFS.

### Estimation

Using the training data sets, we estimated the one-dimensional SFS for YRI and Maasai Mara, as well as the two-dimensional SFS for CEU/YRI and Shangani/Maasai Mara. We ran `winsfs` for 500 epochs using a fixed number of blocks *B* = 500 and window sizes *W* ∈ {100, 250, 500}.

We will focus on the setting with window size *W* = 100. For convenience, we introduce the notation `winsfs`_100_ to refer to `winsfs` with hyperparameter settings *B* = 500, *W* = 100. We return to the topic of hyperparameter settings below.

To compare, we ran `realSFS` using default settings, except allowing it to run for a maximum of 500 epochs rather than the default 100. We will still take the 100 epochs cut-off to mark convergence, if it has not occured by other criteria before then, but results past 100 will be shown in places.

In each case, we evaluated the full log-likelihood (eq. (8)) of the estimates after each epoch on both the training and test data sets. In addition, we computed various summary statistics from the estimates after each epoch. For details, see supplementary text S2.

### One-dimensional SFS

Main results for the one-dimensional estimates are shown in figure 2.

**Figure 2:**
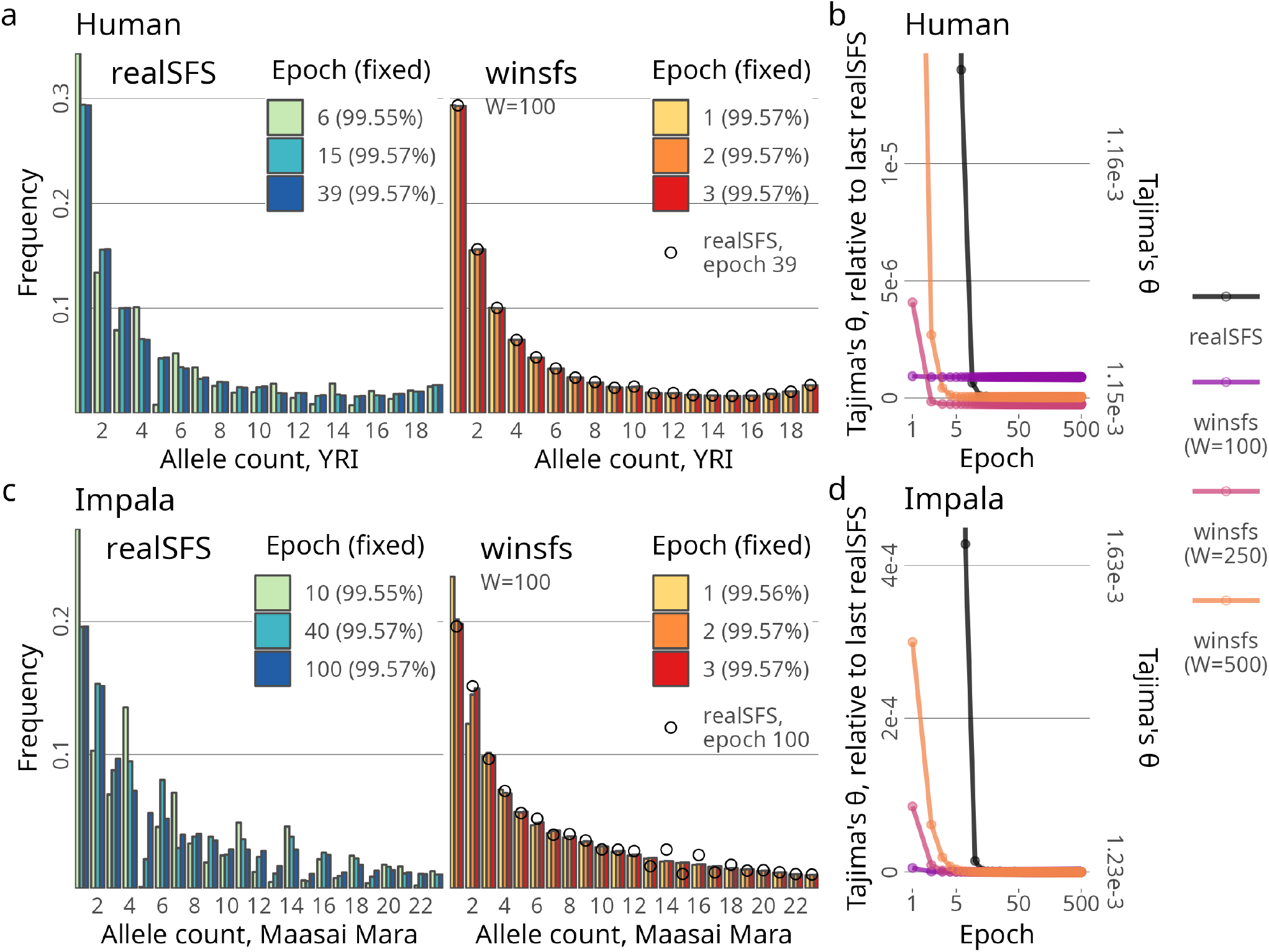
One-dimensional SFS estimation. (a): YRI SFS estimates from `realSFS` and `winsfs`_100_ after various epochs. Only variable sites are shown, proportion of fixed sites is shown in the legend. The final `realSFS` estimate is overlaid with dots on the winsfs plot for comparison. (b): YRI Tajima’s θ estimates calculated from `realSFS` and `winsfs` over epochs. (c): Maasai Mara SFS estimates from `realSFS` and `winsfs`_100_ after various epochs. Only variable sites are shown, proportion of fixed sites is shown in the legend. The final `realSFS` estimate is overlaid with dots on the winsfs plot for comparison. (d): Maasai Mara Tajima’s θ estimates calculated from `realSFS` and `winsfs` over epochs.

For the human YRI population, we find that a single epoch of `winsfs`_100_ produces an estimate of the SFS that is visually indistinguishable from the converged estimate of `realSFS` at 39 epochs (figure 2a). Train and test set log-likelihoods (supplementary figure 3) confirm that the likelihood at this point is only very marginally lower for `winsfs`_100_ than the last `realSFS`. By increasing the window size to 250 or 500, we get test log-likelihood values equal to or above those achieved by `realSFS`, and still within the first 5 epochs.

As an example of a summary statistic derived from the one-dimensional SFS, figure 2b shows that `winsfs`_100_ finds an estimate of Tajima’s *θ* that is very near to the final `realSFS`, with a difference on the order of 1 · 10^−6^. Increasing the window size removes this difference at the cost of a few more epochs.

In the case of Maasai Mara, `realSFS` runs for the 500 epochs, so we take epoch 100 to mark convergence. On this data, `winsfs`_100_ requires two epochs to give a good estimate of the SFS, as shown in figure 2c. Some subtle differences relative to the `realSFS` results remain, however, especially at the middle frequencies: the `realSFS` estimate exhibits a ‘wobble’ such that even bins are consistently higher than odd bins. Such a pattern is not biologically plausible, and is not seen in the `winsfs` spectrum.

Supplementary figure 4 shows train and test log-likelihood data for Maasai Mara, which again support the conclusions drawn from looking at the estimates themselves. In theory, we expect that the test log-likelihood should be adversely impacted by the `realSFS` ‘wobble’ pattern. In practice, however, with more than 99.5 % fixed sites, the fixed end of the spectrum dominate the likelihood to the extent that the effect is not visible. We return to this point below.

Finally, Figure 2d shows that Tajima’s *θ* is likewise well-estimated by one or two epochs of `winsfs`_100_ on the impala data.

### Two-dimensional SFS

Overall results for the joint spectra are seen in figure 3.

**Figure 3:**
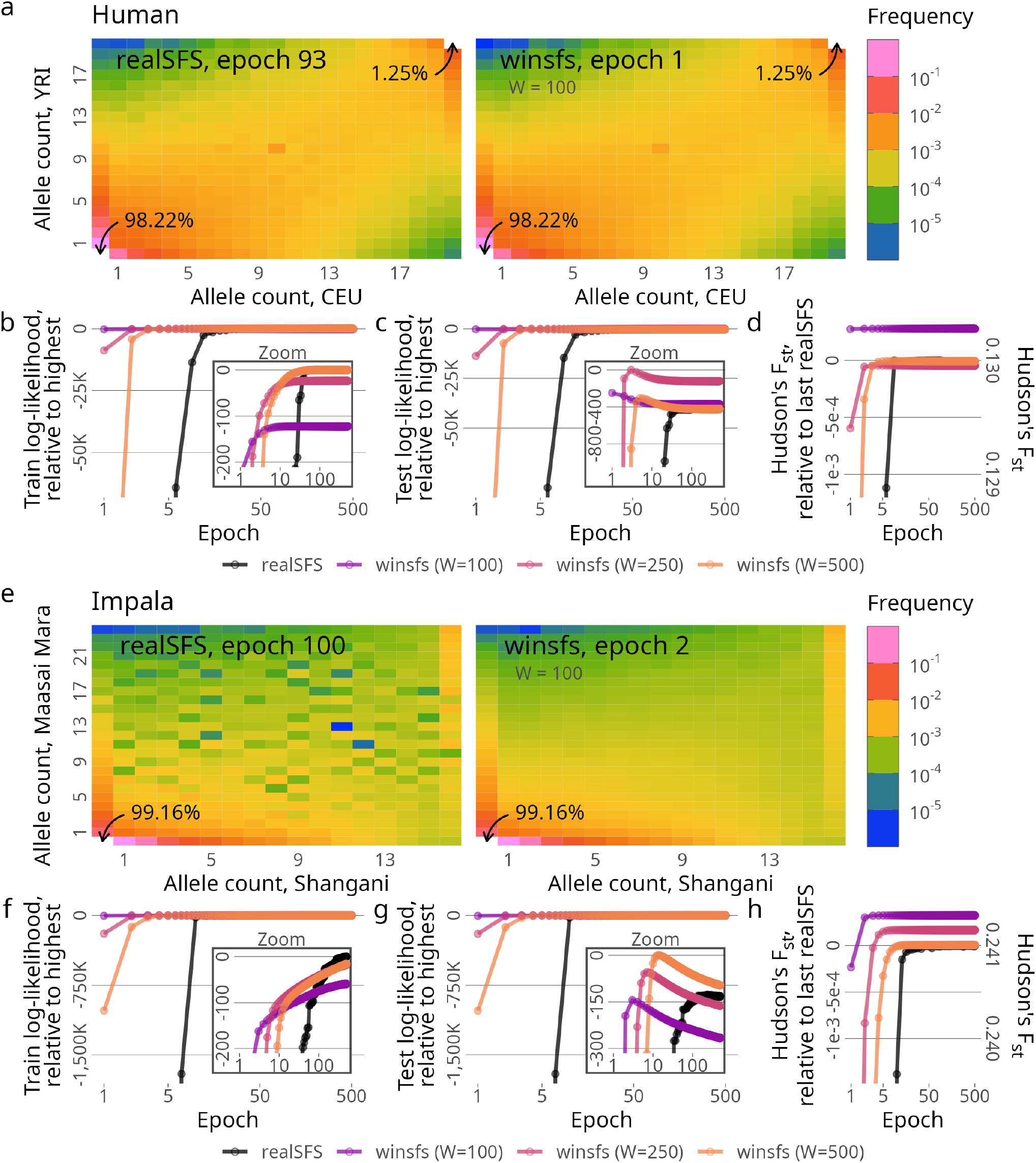
Two-dimensional SFS estimation. (a): CEU/YRI SFS estimates from `realSFS` after 93 epochs (converged) and from `winsfs`_100_ after a single epoch. Fixed sites not shown for scale, total proportion indicated by arrows. (b), (c):CEU/YRI SFS train and test log-likelihood over epochs for `realSFS` and `winsfs`. (d): CEU/YRI Hudson’s F_st_ estimates calculated from `realSFS` and `winsfs` over epochs (e): Shangani/Maasai Mara SFS estimates from `realSFS` after 100 epochs (converged) and from `winsfs`_100_ after a single epoch. Fixed reference sites not shown for scale, proportions indicated by arrows. (f), (g): Shangani/Maasai Mara SFS train and test log-likelihood over epochs for `realSFS` and `winsfs`. (h): Shangani/Maasai Mara Hudson’s F_st_ estimates calculated from `realSFS` and `winsfs` over epochs.

On the human data, `winsfs`_100_ takes a single epoch for an estimate of the SFS that is near-identical to `realSFS` at convergence after 93 epochs. Looking at the log-likelihood results, it is notable that while `realSFS` does better than `winsfs` when evaluated on the training data (figure 3b), the picture is reversed when evaluated on the test data (figure 3c). In fact, all `winsfs` hyperparameter settings achieved better test log-likelihood values in the first 10 epochs than achieved by `realSFS` at convergence. This is likely caused by a faint ‘checkerboard’ pattern in the `realSFS` estimate due to overfitting, as we expect the spectrum to be smooth. We note that both `realSFS` and `winsfs` preserve an excess of sites where all individuals are heterozygous, corresponding to the peak in the centre of the spectrum. This is a known issue with this data set [26], likely caused by paralogs in the mapping process. It is an artefact which can be removed by filtering the data before SAF calculation, which we have not done here. Given this choice, it is to be expected that this peak remains.

In two dimensions, we compute both Hudson’s F_st_ (figure 3d) and the *f*_2_-statistic (supplementary figure 5) from SFS estimates after all epochs, and we note similar patterns for these as we have seen before: one epoch of `winsfs`_100_ gives an estimate of the summary statistic that is almost identical to the final `realSFS` estimate.

For the impalas, `winsfs`_100_ requires two epochs for a good estimate of the spectrum, while `realSFS` again does not report convergence within the first 100. What is immediately striking about the impala results, however, is that the checkerboard pattern is very pronounced for `realSFS`, and again absent for `winsfs` (figure 3e). The problem for `realSFS` is likely exacerbated by two factors: first, the sequencing depth is lower, increasing the uncertainty; second, the relatively high divergence of the impala populations push most of the mass in the spectrum towards the edges. Together, this means that very little information is available for most of the estimated parameters. It appears that `realSFS` therefore ends up overfitting to the particularities of the training data at these bins.

This is also reflected in the difference between train and test log-likelihood (figures 3f and 3g). Like in the case of the human data, the SFS estimated by `winsfs` performs better on the test data compared to `realSFS`, while `realSFS` performs the based on the training data. On the test data, all `winsfs` settings again reach log-likelihood values comparable to or better than `realSFS` in few epochs. However, the differences between `realSFS` and `winsfs` remain relatively small in terms of log-likelihood, even on the test set. This is somewhat surprising, given the marked checkerboarding in the spectrum itself. Again, we attribute this to the fact that the log-likelihood is dominated by all the mass lying in or around the zero-zero bin. We expect, therefore, that methods that rely on the ‘interior’ of the SFS should do better when using `winsfs`, compared to `realSFS`.

Before turning to test this prediction, we briefly note that F_st_ (figure 3h) and the *f*_2_-statistic (supplementary figure 5) are also adequately estimated for the impalas by `winsfs`_100_ in one epoch.

### Demographic inference

All the SFS-derived summary statistics considered so far are heavily influenced by the bins with the fixed allele bins (that is, count 0 or 2*N*_*k*_ in all populations), or they are sums of alternating frequency bins. In either case, this serves to mask issues with checkerboard areas of the SFS in the lower-frequency bins. However, this will not be the case for downstream methods that rely on the shape of the spectrum in more detail.

To illustrate, we present a small case-study of inferring the demographic history of the impala populations using the *∂;a∂;i* [5] software with the estimated impala spectra shown in figure 3e, though folded due to the lack of an outgroup for proper polarisation. Briefly, based on an estimated SFS and a user-specified demographic model, *∂;a∂;i* fits a model SFS based on the demographic parameters so as to maximise the likelihood of these parameters.

Our approach was to fit a simple demographic model for the Shangani and Maasai Mara populations, and then gradually add parameters to the model as required based on the residuals of the input and model spectra. We take this to be representative of a typical workflow for demographic inference.

For each successive demographic model [27], we ran *∂;a∂;i* on the folded spectra by performing 100 independent optimisation runs from random starting parameters, and checking for convergence by requiring the top three results to be within five likelihoods units of each other. If the optimisation did not converge, we did additional optimisation runs until either they converged or 500 independent runs were reached without likelihood convergence. In that case, we inspected the results for the top runs, to assess whether they were reliably reaching similar estimates and likelihoods. Results are shown in figure 4.

**Figure 4:**
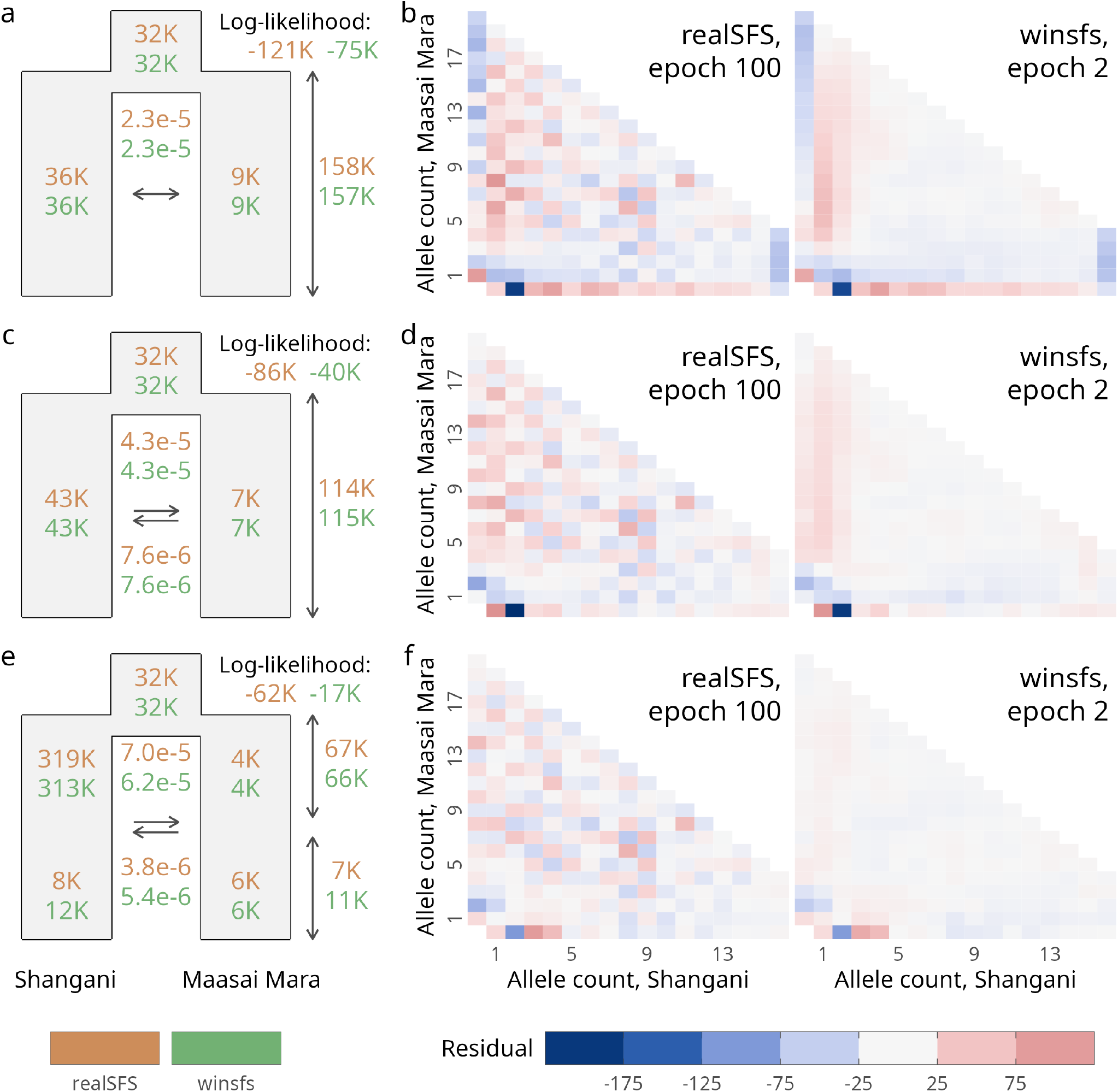
Demographic inference results. Each row corresponds to a demographic model fitted using ∂a∂i. On the left, a schematic of the model is shown including parameter estimates using SFS estimates from `realSFS` after 100 epochs or from `winsfs`_100_ after two epochs. Time is given in years, population sizes in number of individuals, and migration rates is per chromosome per generation. All parameters were scaled assuming a mutation rate of 1.41 · 10^−8^ per site per generation and a generation time of 5.7 years. On the right, the residuals of the SFS fitted by ∂a∂i. Note that ∂a∂i folds the input SFS, hence the residuals are likewise folded. The fixed category is omitted to avoid distorting the scale. (a), (b): Model with symmetric migration and constant population size. (c), (d): Model with asymmetric migration and constant population size. (e), (f): Model with asymmetric migration and a single, instantaneous population size change.

The first, basic model assumes that the populations have had constant populations sizes and a symmetric migration rate since diverging. The parameter estimates based on `realSFS` and `winsfs` are similar, though the `winsfs` model fit has significantly higher log-likelihood (figure 4a). However, when inspecting the residuals in figure 4b, the `realSFS` residuals suffer from a heavy checkerboard pattern, making it hard to distinguish noise from model misspecification. In contrast, the `winsfs` residuals clearly show areas of the spectrum where the model poorly fits the data.

In particular, the residuals along the very edge of the spectrum suggest that a symmetric migration rate is not appropriate. Therefore, we fit a second model with asymmetric migration (figure 4c) Now *∂a∂i* finds migration rates from Shangani to Maasai Mara an order of magnitude higher than *vice versa*. The results for `winsfs` (figure 4d) show improved residuals, while the `realSFS` residuals remain hard to interpret.

Finally, an area of positive residuals in the fixed and rare-variant end of the Shangani spectrum suggests that this population has recently undergone a significant bottleneck. Therefore, the third model allows for an instantaneous size change in each of the impala populations (figure 4e). At this point, the `winsfs` residuals (figure 4f) are negligible, suggesting that no more parameters should be added to the model. Once again, though, the `realSFS` residuals leave us uncertain whether further model extensions are required.

When looking at the final model fits, the *∂;a∂;i* parameter estimates from `realSFS` and `winsfs` also start to differ slightly. In several instances, estimates disagree by about 50 %, and the log-likelihood remains much higher for `winsfs`, with a difference of 45 000 log-likelihood units to `realSFS`. In addition, we confirmed that the log-likelihood of the original test data set given the SFS fitted by *∂;a∂;i* is higher for `winsfs` (−8.08 · 10^8^) than for `realSFS` (−8.38 · 10^8^). We stress, however, that we would have likely never found the appropriate model without using `winsfs`, since the interpretation of the `realSFS` results is difficult. In relation to this point, we note that the final model results in considerably different estimates for parameters of biological interest, such as split times and recent population sizes, relative to the initial model. We also find that the last model is supported by the literature: previous genetic and fossil evidence suggests extant common impala populations derive from a refugia in Southern Africa that subsequently colonised East Africa in the middle-to-late Pleistocene [28–30]. This is broadly consistent with the estimated split time, and the reduction in population size in East African populations as they colonised the new habitat. The difference in effective population size between the southern Shangani population and the eastern Maasai Mara was previously also found using microsatellite data [28].

### Simulations

To validate these findings in conditions with a known SFS, we ran simulations using `msprime` [31] and `tskit` [32]. Briefly, we simulated two populations, which we simply refer to as A and B. Populations A and B diverged 10 000 generations ago and both have effective populations sizes of 10 000 individuals, except for a period of 1000 generations after the split, during which time B went through a bottleneck of size 1000. We simulated 22 independent chromosomes of 10 Mb for a total genome size of 220 Mb, using a mutation rate of 2.5 · 10^−8^ and a uniform recombination rate of 1 · 10^−8^. To explore the consequences of varying sample sizes, we sampled 5, 10, or 20 individuals from the two populations. For each of these three scenarios, we calculated the true SFS from the resulting genotypes (shown in supplementary figure 6).

Using the true genotypes as input, we simulated the effects of NGS sequencing with error for both the variable and invariable sites. At every position in the genome, including the monomorphic sites, we sample *D* ∼ Poisson(*λ*) bases and introduce errors with a constant rate of *ε* = 0.002 independently for each base. We calculate genotype likelihoods according to the GATK model outlined in equations (1) to (3) and output GLF files. Using these, we create SAF files for A and B with no further filtering using `ANGSD`. The mean depth *λ* is set to either 2, 4, or 8 to investigate the performance of `winsfs` at difference sequencing depths. This results in a grid of 3 × 3 simulated NGS data sets with three different sample sizes and three different mean depth values.

From the simulated SAF files, we ran `winsfs` and `realSFS` as above to generate the two-dimensional SFS, except for a maximum of 100 epochs. For each method and each epoch *e* until convergence, we calculated the log-likelihood for the corresponding SFS 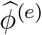,

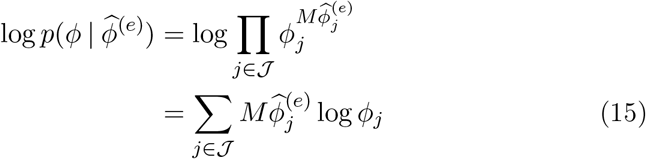

where *ϕ* is the observed true SFS and *M* is the total number of sites. Figure 5 shows how the log-likelihood evolves over epochs for `winsfs` (*W* ∈ {100, 250, 500}) and `realSFS` for sample sizes *N*_*k*_ ∈ {5, 10, 20} and simulated mean depths *λ* ∈ {2, 4, 8}. We observe that at a mean depth of 2, `winsfs`_100_ outperforms `realSFS` by a significant margin both in terms of speed and the final log-likelihood. At mean depth 4, the `winsfs` remains much faster and still achieves meaningfully better log-likelihoods, especially at higher sample sizes. Finally, at mean depth 8, `winsfs`_100_ still converges 5–10 times faster than `realSFS` (measured in epochs), but the methods provide estimates of similar quality.

**Figure 5:**
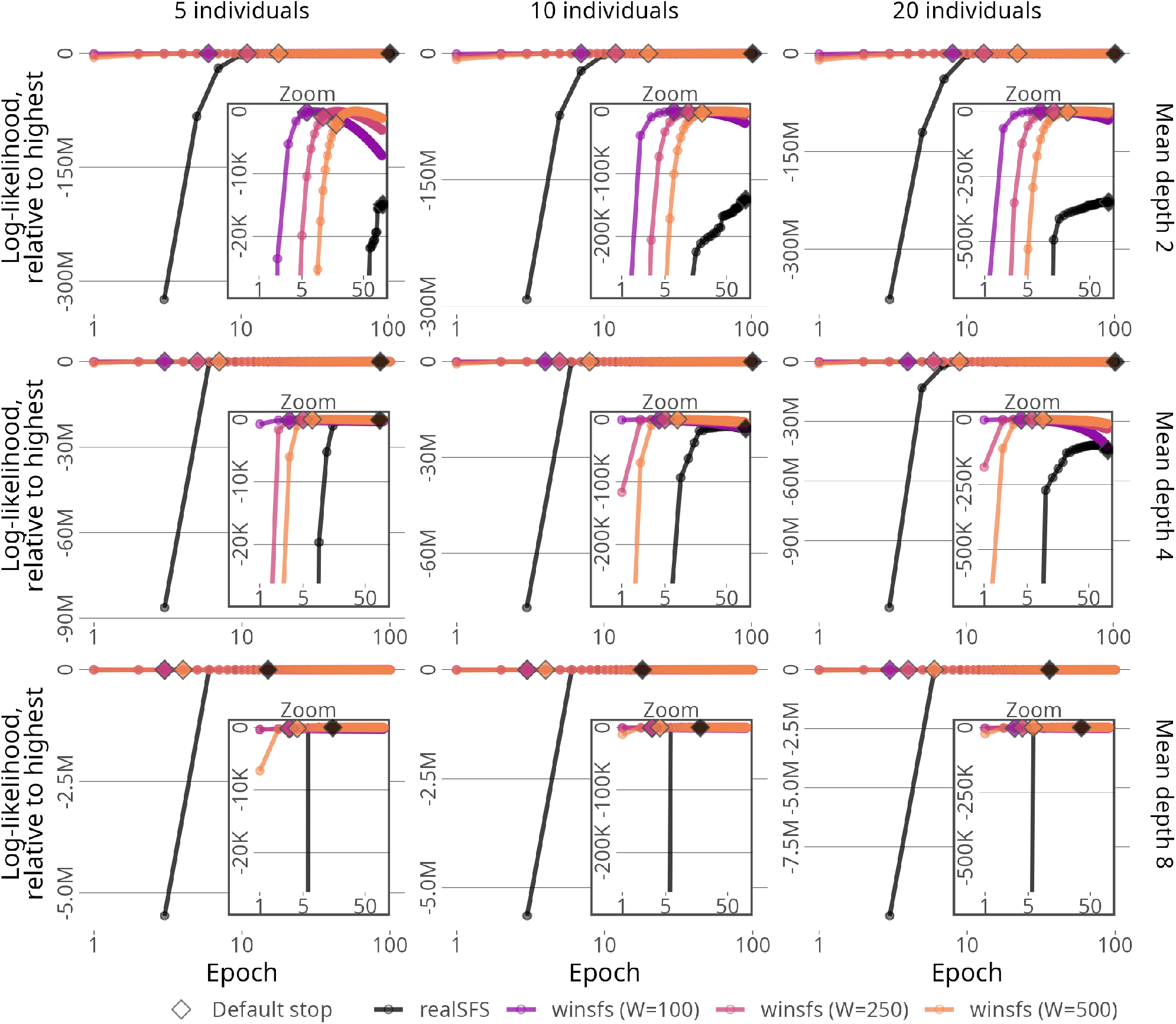
Log-likelihood over epochs of the true observed SFS given the two-dimensional SFS estimated by `winsfs` (*W* ∈; {100, 250, 500}) and `realSFS`. Different simulated scenarios (mean depth 2, 4, or 8; sample size 5, 10, or 20) shown. For each method, the epoch at which the default stopping criterion is triggered is shown. Note that the y-scale varies across sample sizes and depths in order to show the full range of data (main plot) and the difference between `realSFS` and `winsfs` (zoom plot). For each column of plots, corresponding to a simulated sample size, the *y*-scale in the zoom plot is held constant to allow for comparison across depths.

The estimated spectra for `realSFS` and `winsfs`_100_ at their default stopping points are shown in supplementary figure 7 and supplementary figure 8 and respectively. These confirm that the spectra on the whole are well-estimated by `winsfs`_100_ as compared to the true SFS (supplementary figure 6). Moreover, we again observe that `realSFS` introduces a checkerboard pattern in the low-information part of the spectrum at 2 x–4 x, which is not present in the true spectrum, and which is not inferred by `winsfs`. The pattern is more pronounced at higher sample sizes. This supports the hypothesis that `realSFS` tends to overfit in situations where many parameters must be inferred with little information.

### Peak simulations

The averaging of block estimates in the window EM algorithm appears to induce a certain ‘smoothing’ of the spectrum at low depth. This smoothing effect is implicit in the sense of being nowhere explicitly modelled, and each parameter is estimated independently. Nevertheless, this observation may give rise to a concern that `winsfs`, unlike the maximum likelihood estimate from `realSFS`, might remove true abrupt peaks in the SFS.

To investigate, we modified the demographic simulation with sample size 20 described above in the following way. In each of seven arbitrarily chosen bins near to the centre of the SFS, we artificially spiked 10 000 counts into the true spectrum after running the demographic simulations (supplementary figure 9). This represents a 30–40-fold increase relative to the original count and the neighbouring cells. Based on this altered spectrum, we simulated sequencing data for depth 2 x, 4 x, and 8 x, created SAF files, and ran `realSFS` and `winsfs`_100_ as before. The residuals of the `realSFS` and `winsfs` estimates are shown in supplementary figure 10 and supplementary figure 11, respectively. In this fairly extreme scenario, the spectra inferred by both `winsfs` and `realSFS` appear to have a small but noticeable downwards bias in the peak region at 2 x and 4 x. However, compared to `realSFS`, `winsfs` has smaller residuals in all scenarios, and the apparent bias is inversely correlated with depth. These results confirm that usage the window EM algorithm does not lead to excess flattening of SFS peaks compared with the maximum likelihood estimate from the standard EM algorithm.

### Hyperparameters

The window EM algorithm requires hyperparameter settings for *B* and *W*. Moreover, it requires a choice of stopping criterion. For ease of use, the `winsfs` software ships with defaults for these settings, and we briefly describe these.

We expect that the choice of *B* is less important than the term *W/B*, which governs the fraction of data that is directly considered in any one update step. Having analysed input data varying in size from 220 Mb (simulations) to 1.17 Gb (human data), we find that fixing *B* = 500 works fine as a default across a wide range of input sizes. Therefore, the more interesting question is how to set the window size. In theory, there should be a trade-off between speed of convergence and accuracy of results, where lower window size favours the former and higher window size the latter. However, in practice, based on our results, we have not seen evidence that using *W* = 500 over *W* = 100 leads to significantly better inference. On the other hand, the lower window size has significantly faster convergence. Based on this, we feel that window size of 100 makes for the best general default. By default, the `winsfs` software uses *B* = 500 blocks and a window size *W* = 100.

As for stopping, `winsfs` implements the criterion based differences *δ* In 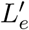 (eq. (14)) over successive epochs. Based on the initial analysis of the human and impala data, we chose *δ* = 10^*−*4^ (see supplementary figure 12) as the default value and used the simulations to validate this choice. Figure 5 shows the point at which stopping occurs, which is generally around the maximum log-likelihood as desired.

### Streaming

In the main usage mode, pre-calculated SAF likelihoods are read into RAM, as in `realSFS`. However, it is also possible to run `winsfs` while keeping the data on disk and streaming through the intersecting sites in the SAF files. We refer to this as ‘streaming mode’.

Since the window EM algorithm requires randomly shuffling the input data, a preparation step is required in which SAF likelihoods are (jointly) shuffled into a new file. We wish to avoid loading the data into RAM in order to perform a shuffle, and we also do not want multiple intermediate writes to disk. To our knowledge, it is not possible to perform a true shuffle of the input data within these constraints. Instead, since we are only interested in shuffling for the purposes of breaking up blocks of LD, we perform a pseudo-shuffle according to the following scheme. We pre-allocate a file with space for exactly *M* intersecting sites in the input data. This file is then split into *S* contiguous sections of roughly equal size, and we then assign input site with index *m* ∈ {1, …, *M*} to position └ (*m* + 1)*/S*┘ + 1 in section (*m* + 1) % *S* + 1, where % is the remainder operation. That is, the first *S* sites in the input end up in the first positions of each section, and the next *S* sites in the input end up in the second positions of each section, and so on. This operation can be performed with constant memory, without intermediate writes to disk, and has the benefit of being reversible.

After preparing the pseudo-shuffled file, `winsfs` can be run exactly as in the main mode. To confirm that this pseudo-shuffle is sufficient for the purposes of the window EM algorithm, we ran 10 epochs of `winsfs` in streaming mode for the impala and human data sets in both one and two dimensions. After each epoch, we calculated the log-likelihood of the resulting SFS and compared them to the log-likelihood obtained by running in main mode above. The results are shown in supplementary figure 13 and show that streaming mode yields comparable results to the main, in-RAM usage: the likelihood differs slightly, but is neither systematically better or worse.

### Benchmark

To assess its performance characteristics, we benchmarked `winsfs` in both the main mode and streaming mode as well as `realSFS` on the impala data. For each of the three, we ran estimation until convergence, as well as until various epochs before then, collecting benchmark results using Snakemake [33]. Both `realSFS` and `winsfs` were given 20 cores. Results are shown in figure 6. In terms of run-time, we find that running `winsfs` in RAM is significantly faster than `realSFS` (figure 6a). This is true in part because `winsfs` requires fewer epochs, but also since `winsfs` runs faster than `realSFS` epoch-by-epoch. As expected, when switching `winsfs` to streaming mode, run-time suffers as epochs increase. However, taking the number of epochs required for convergence into account, streaming `winsfs` remains competitive with `realSFS`, even when including the initial overhead to shuffle SAF likelihoods on disk.

**Figure 6:**
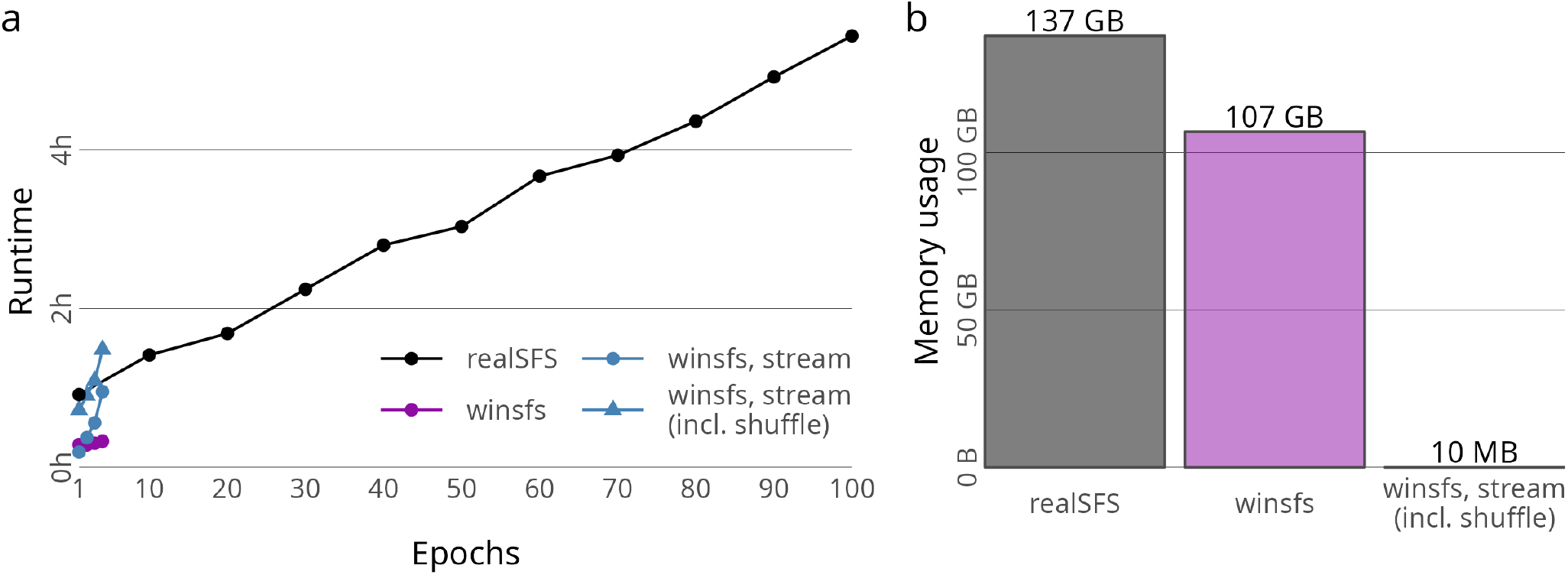
Computational resource usage of `winsfs` and `realSFS` for the joint estimation of the Shangani and Maasai Mara impala populations `winsfs` can be run while loading input data into RAM, or streaming through it on disk. In the latter case, data must be shuffled on disk before hand. (a): Runtime required with 20 threads for various numbers of epochs. Results for `winsfs` are shown for in-memory usage and streaming mode. For streaming modes, times are given with and without the extra time taken to shuffle data on disk before running. (b): Peak memory usage (maximum resident set size).

Looking at memory consumption, streaming `winsfs` has a trivial peak memory usage of 10 MB, including the initial pseudo-shuffle. In comparison, when reading data into RAM, `realSFS` and `winsfs` require 137 GB and 107 GB, respectively, even on the fairly small impala data set.

The benchmarking results for the one-dimensional Maasai Mara estimation are shown in supplementary figure 14 and support similar conclusions.

## 4 Discussion

We have presented the window EM algorithm for inferring the SFS from low-depth data, as well as the `winsfs` implementation of this algorithm. The window EM algorithm updates SFS estimates in smaller blocks of sites, and averages these block estimates in larger windows. We have argued that this approach has three related advantages relative to current methods. First, by updating more often, convergence happens one to two orders of magnitude faster. Due to the window averaging, this improvement in convergence times does not occur at the cost of stability. Second, due to the fast convergence, it is feasible to run the window EM algorithm out of memory. This brings the memory requirements of the algorithm from hundreds of gigabytes of RAM to virtually nothing. Third, by optimising over different subsets of the data in each iteration, the algorithm is prevented from overfitting to the input data. In practice, this means we get biologically more plausible spectra.

On this last point, it is worth emphasising that while `winsfs` appears to have the effect of smoothing the spectrum in a beneficial way, this smoothing effect is entirely implicit. That is, it is nowhere explicitly modelled that each estimated bin should be similar to neighbouring bins to avoid checkerboard patterns. Rather, the apparent smoothing emerges because `winsfs` mitigates some of the issues with overfitting that may otherwise manifest as a checkerboard pattern. As shown in the simulations, `winsfs` does not remove true peaks in the SFS. In the broader setting of stochastic optimization, window EM is in this way related to forms of Polyak-Ruppert iterate averaging schemes as used in stochastic gradient methods [34, 35], variants of which have also been shown to control variance and induce regularisation [36, 37], similar to what we have observed here.

Within the EM literature, window EM is *prima facie* quite similar in spirit to other versions of the stochastic EM algorithm [38–42]. They too work on smaller blocks, and seek some way of controlling stability in how the block estimate 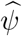 is incorporated in the overall estimate 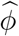. Typically, this involves an update of the form 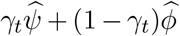 for some weight *γ*_*t*_ decaying as a function of iteration *t*. During initial experimentation, we empirically found that such methods tended to increase the noise in the spectrum, rather than reduce it. This problem likely arises because estimating the multidimensional SFS requires estimating many parameters for which very little information is available in any one batch. Therefore, by having an update step involving only the current estimate and a single, small batch of sites, significant noise is introduced in the low-density part of the spectrum. In contrast, the window EM approach still optimises over smaller batches for speed, but actually considers large amounts of data in the update step by summing the entire window of batch estimates, thereby decreasing the noise.

For SFS inference specifically, prior work exists to improve estimation for low-depth sequencing data. For example, it has been proposed to ‘band’ SAF likelihoods to make estimation scale better in the number of sampled individuals [23, 43]. Briefly, the idea is that at each site, all the mass in the SAF likelihood tends to be concentrated in a small band around the most likely sample frequency, and downstream inference can be adequately carried out by only propagating this band and setting all others to zero. By doing so, run-time and RAM can be saved by simply ignoring all the zero bins outside the chosen band. We note that such ideas are orthogonal to the work presented here, since they are concerned with the representation of the input data, and thereby indirectly modify all downstream optimisation methods. Future work on `winsfs` may involve the ability to run from banded SAF likelihoods. This will be important with large sample sizes, in the hundreds of individuals.

Others have focused on the implementation details of the EM algorithm, for instance using GPU acceleration [44]. Such efforts still have the typical high memory requirements, and do not address the overfitting displayed by the standard EM algorithm. Moreover, we find that the presented algorithmic improvements, combined with an efficient implementation, serve to make `winsfs` more than competitive with such efforts in terms of runtime. Indeed, with `winsfs` converging in-memory in less than an hour on genome-scale data, runtime is no longer a significant bottleneck for SFS estimation.

We emphasise, however, that the window EM algorithm and `winsfs` are unlikely to yield any meaningful benefits with sequencing data at above around 10 x–12 x coverage. With such data, better inference of the SFS will be obtained by estimation directly from genotype calls with appropriate filters. Nevertheless, efficient and robust methods remain important for low-coverage data. This is partly because low-coverage data may sometimes be the only option, for example when working with ancient DNA. Also, such methods allow intentionally sequencing at lower coverage, decreasing the sequencing cost per individual.

In addition, we do not expect `winsfs` to perform better than `realSFS` when data is not available for many sites (e.g <100 Mb) due to the fact that `winsfs` only uses parts of the available data directly in the final estimation.

Finally, improvements in the SFS estimates by `winsfs` are unlikely to be significant for simple summary statistics like *θ*, F_st_, or *f*-statistics. For such purposes, `winsfs` simply produces results similar to `realSFS`, although much faster. However, as the number of dimensions and samples increase, and as sequencing depth decreases, overfitting will start to influence the low-frequency bins of the spectrum. Where this information is used downstream, `winsfs` will lead to better and more interpretable results, and can potentially help solve commonly known biases in parameter estimates arising from model misspecification [45]. We have seen this in the *∂a∂i* case study, but we believe the same would be true of other popular demographic inference frameworks including fastsimcoal [6, 46], moments [47], and momi [48]. It may also be significant for other methods for complex inference from the multidimensional spectrum, including inference of fitness effects using fit*∂a∂i* [49, 50] or introgression using D_FS_ [51], though we have not explored these methods.

## 5 Code and data availability

The human data analysed is part of the 1000 Genomes [25] phase 3 low depth sequencing data. Alignments have been made available by the 1000G project and can be accessed at ftp.1000genomes.ebi.ac.uk/vol1/ftp/phase3/. The impala data has been made available via the SRA with accession `PRJNA862915`. Analysis and plotting code, as well as the cleaned data corresponding to the final results, is available at github.com/malthesr/window and the `winsfs` software itself at github.com/malthesr/winsfs.

## 6 Acknowledgements

AA, CW, and MSR are supported by the Independent Research Fund Denmark (grant numbers: 8021-00360B, 0135-00211B) and the University of Copenhagen through the Data+ initiative. GGE is supported by the Independent Research Fund Denmark (grant number: 8049-00098B). TSK is funded by a Carlsberg Foundation Young Researcher Fellowship awarded by the Carlsberg Foundation in 2019 (CF19-0712).

## References

1. Korneliussen, T. S., Moltke, I., Albrechtsen, A. & Nielsen, R. & Nielsen, R. Calculation of Tajima’s D and Other Neutrality Test Statistics from Low Depth Next-Generation Sequencing Data. BMC Bioinformatics 14 (2013).

2. Bhatia, G., Patterson, N., Sankararaman, S. & Price, A. L. Estimating and Interpreting F<sub>ST</sub>: The Impact of Rare Variants. Genome Research 23, 1514–1521 (2013).

3. Peter, B. M. Admixture, Population Structure, and F-Statistics. Genetics 202, 1485–1501 (2016).

4. Marth, G. T., Czabarka, E., Murvai, J. & Sherry, S. T. The Allele Frequency Spectrum in Genome-Wide Human Variation Data Reveals Signals of Differential Demographic History in Three Large World Populations. Genetics 166, 351–372 (2004).

5. Gutenkunst, R. N., Hernandez, R. D., Williamson, S. H. & Bustamante, C. D. Inferring the Joint Demographic History of Multiple Populations from Multidimensional SNP Frequency Data. PLoS Genetics 5 (ed McVean, G.) e1000695 (2009).

6. Excoffier, L., Dupanloup, I., Huerta-Sánchez, E., Sousa, V. C. & Foll, M. Robust Demographic Inference from Genomic and SNP Data. PLoS Genetics 9 (ed Akey, J. M.) e1003905 (2013).

7. Tajima, F. Statistical Method for Testing the Neutral Mutation Hypothesis by DNA Polymorphism. Genetics 123, 585–595 (1989).

8. Fay, J. C. & Wu, C.-I. Hitchhiking Under Positive Darwinian Selection. Genetics 155, 1405–1413 (2000).

9. Nielsen, R. et al. A Scan for Positively Selected Genes in the Genomes of Humans and Chimpanzees. PLoS Biology 3 (ed Tyler-Smith, C.) e170 (2005).

10. Nielsen, R., Paul, J. S., Albrechtsen, A. & Song, Y. S. Genotype and SNP Calling from Next-Generation Sequencing Data. Nature Reviews Genetics 12, 443–451 (2011).

11. Han, E., Sinsheimer, J. S. & Novembre, J. Characterizing Bias in Population Genetic Inferences from Low-Coverage Sequencing Data. Molecular Biology and Evolution 31, 723–735 (2013).

12. Olalde, I. et al. The Genomic History of the Iberian Peninsula Over the Past 8000 Years. Science 363, 1230–1234 (2019).

13. Margaryan, A. et al. Population Genomics of the Viking World. Nature 585, 390–396 (2020).

14. Van der Valk, T. et al. Million-Year-Old DNA Sheds Light on the Genomic History of Mammoths. Nature 591, 265–269 (2021).

15. Lou, R. N., Jacobs, A., Wilder, A. P. & Therkildsen, N. O. A Beginner’s Guide to Low-Coverage Whole Genome Sequencing for Population Genomics. Molecular Ecology 30, 5966–5993 (Aug. 2021).

16. Li, H. A Statistical Framework for SNP Calling, Mutation Discovery, Association Mapping and Population Genetical Parameter Estimation from Sequencing Data. Bioinformatics 27, 2987–2993 (2011).

17. Sánchez-Barreiro, F. et al. Historical Population Declines Prompted Significant Genomic Erosion in the Northern and Southern White Rhinoceros (Ceratotherium simum). Molecular Ecology 30, 6355–6369 (2021).

18. Li, H. et al. The Sequence Alignment/Map format and SAMtools. Bioinformatics 25, 2078–2079 (2009).

19. Danecek, P. et al. Twelve Years of SAMtools and BCFtools. GigaScience 10 (2021).

20. McKenna, A. et al. The Genome Analysis Toolkit: A MapReduce Framework for Analyzing Next-Generation DNA Sequencing Data. Genome Research 20, 1297–1303 (2010).

21. Korneliussen, T. S., Albrechtsen, A. & Nielsen, R. ANGSD: Analysis of Next Generation Sequencing Data. BMC Bioinformatics 15 (2014).

22. Nielsen, R., Korneliussen, T., Albrechtsen, A., Li, Y. & Wang, J. SNP Calling, Genotype Calling, and Sample Allele Frequency Estimation from New-Generation Sequencing Data. PLoS ONE 7 (ed Awadalla, P.) e37558 (2012).

23. Han, E., Sinsheimer, J. S. & Novembre, J. Fast and accurate site frequency spectrum estimation from low coverage sequence data. Bioinformatics 31, 720–727 (2014).

24. Varadhan, R. & Roland, C. Simple and Globally Convergent Methods for Accelerating the Convergence of Any EM Algorithm. Scandinavian Journal of Statistics 35, 335–353 (2008).

25. The 1000 Genomes Project Consortium. A Global Reference for Human Genetic Variation. Nature 526, 68–74 (2015).

26. Meisner, J. & Albrechtsen, A. Testing for Hardy–Weinberg Equilibrium in Structured Populations Using Genotype or Low-Depth Next Generation Sequencing Data. Molecular Ecology Resources 19, 1144–1152 (June 2019).

27. Portik, D. M. et al. Evaluating mechanisms of diversification in a Guineo-Congolian tropical forest frog using demographic model selection. Molecular Ecology 26, 5245–5263 (Aug. 2017).

28. Lorenzen, E. D., Arctander, P. & Siegismund, H. R. Regional Genetic Structuring and Evolutionary History of the Impala Aepyceros melampus. Journal of Heredity 97, 119–132 (2006).

29. Lorenzen, E. D., Heller, R. & Siegismund, H. R. Comparative phylogeography of African savannah ungulates. Molecular Ecology 21, 3656–3670 (2012).

30. Faith, J. T., Tryon, C. A., Peppe, D. J., Beverly, E. J. & Blegen, N. Biogeographic and Evolutionary Implications of an Extinct Late Pleistocene Impala from the Lake Victoria Basin, Kenya. Journal of Mammalian Evolution 21, 213–222 (2013).

31. Baumdicker, F. et al. Efficient ancestry and mutation simulation with msprime 1.0. Genetics 220 (2021).

32. Kelleher, J., Thornton, K. R., Ashander, J. & Ralph, P. L. Efficient pedigree recording for fast population genetics simulation. PLOS Computational Biology 14, e1006581 (2018).

33. Koster, J. & Rahmann, S. Snakemake–A Scalable Bioinformatics Workflow Engine. Bioinformatics 28, 2520–2522 (Aug. 2012).

34. Ruppert, D. Efficient Estimations from a Slowly Convergent Robbins-Monro Process tech. rep. (Cornell University, 1988).

35. Polyak, B. T. & Juditsky, A. B. Acceleration of Stochastic Approximation by Averaging. SIAM Journal on Control and Optimization 30, 838–855 (1992).

36. Jain, P. et al. A Markov Chain Theory Approach to Characterizing the Minimax Optimality of Stochastic Gradient Descent (for Least Squares) (2018).

37. Neu, G. & Rosasco, L. Iterate Averaging as Regularization for Stochastic Gradient Descent in Conference On Learning Theory (2018), 3222–3242.

38. Neal, R. M. & Hinton, G. E. A View of the EM Algorithm that Justifies Incremental, Sparse, and Other Variants in Learning in Graphical Models 355–368 (Springer Netherlands, 1998).

39. Sato, M.-a. & Ishii, S. On-line EM Algorithm for the Normalized Gaussian Network. Neural Computation 12, 407–432 (2000).

40. Cappé, O. & Moulines, E. On-line Expectation-Maximization Algorithm for Latent Data Models. Journal of the Royal Statistical Society: Series B (Statistical Methodology) 71, 593–613 (2009).

41. Liang, P. & Klein, D. Online EM for Unsupervised Models in Proceedings of Human Language Technologies: The 2009 Annual Conference of the North American Chapter of the Association for Computational Linguistics (2009), 611–619.

42. Chen, J., Zhu, J., Teh, Y. W. & Zhang, T. Stochastic Expectation Maximization with Variance Reduction in Proceedings of the 32nd International Conference on Neural Information Processing Systems 31 (2018), 7967–7977.

43. Mas-Sandoval, A. et al. Fast and Accurate Estimation of Multidimen-sional Site Frequency Spectra from Low-Coverage High-Throughput Sequencing Data. GigaScience 11 (2022).

44. Lu, M., Zhao, J., Luo, Q. & Wang, B. Accelerating Minor Allele Frequency Computation with Graphics Processors in Proceedings of the 1st International Workshop on Big Data, Streams and Heterogeneous Source Mining Algorithms, Systems, Programming Models and Applications - BigMine ‘12 (ACM Press, 2012).

45. Momigliano, P., Florin, A.-B. & Merilä, J. Biases in Demographic Modelling Affect Our Understanding of Recent Divergence. Molecular Biology and Evolution (ed Kim, Y.) (Feb. 2021).

46. Excoffier, L. & Foll, M. fastsimcoal: a Continuous-Time Coalescent Simulator of Genomic Diversity Under Arbitrarily Complex Evolutionary Scenarios. Bioinformatics 27, 1332–1334 (Mar. 2011).

47. Jouganous, J., Long, W., Ragsdale, A. P. & Gravel, S. Inferring the Joint Demographic History of Multiple Populations: Beyond the Diffusion Approximation. Genetics 206, 1549–1567 (July 2017).

48. Kamm, J. A., Terhorst, J. & Song, Y. S. Efficient Computation of the Joint Sample Frequency Spectra for Multiple Populations. Journal of Computational and Graphical Statistics 26, 182–194 (Jan. 2017).

49. Kim, B. Y., Huber, C. D. & Lohmueller, K. E. Inference of the Distribution of Selection Coefficients for New Nonsynonymous Mutations Using Large Samples. Genetics 206, 345–361 (May 2017).

50. Huang, X. et al./person-group>. Inferring Genome-Wide Correlations of Mutation Fitness Effects between Populations. Molecular Biology and Evolution 38 (ed Nielsen, R.) 4588–4602 (May 2021).

51. Martin, S. H. & Amos, W. Signatures of Introgression across the Allele Frequency Spectrum. Molecular Biology and Evolution 38 (ed Harris, K.) 716–726 (Sept. 2020)

