## Supplementary material for "Estimation of site frequency spectra from low-coverage sequencing data using stochastic EM reduces overfitting, runtime, and memory usage"

### Supplementary materials

#### S1 SFS EM algorithm

**General EM** We begin by giving the general proof of convergence for the EM algorithm [1] in the discrete case to establish notation.

For observed data $X$ consisting of $M$ independent data points $X_m$, and discrete latent data $Z_m \in J$, we write the log-likelihood for parameter $\phi$,

$$\begin{aligned} L(\phi) &= \log p(X \mid \phi) \\ &= \sum_{m=1}^{M} \log p(X_m \mid \phi) \\ &= \sum_{m=1}^{M} \log \sum_{j \in J} p(X_m, Z_m = j \mid \phi) \, . \end{aligned} \tag{S.1}$$

For an arbitrary distribution $q_m$ with support $\mathcal{J}$, by Jensen's inequality

$$\begin{aligned} L(\phi) &= \sum_{m=1}^{M} \log \sum_{j \in J} q_m(j) \frac{p(X_m, Z_m = j \mid \phi)}{q_m(j)} \\ &\geq \sum_{m=1}^{M} \sum_{j \in J} q_m(j) \log \frac{p(X_m, Z_m = j \mid \phi)}{q_m(j)} \\ &= \sum_{m=1}^{M} \sum_{j \in J} q_m(j) \log p(X_m \mid \phi) \\ &\quad + \sum_{m=1}^{M} \sum_{j \in J} q_m(j) \log \frac{p(Z_m = j \mid X_m, \phi)}{q_m(j)} \\ &= L(\phi) - \sum_{m=1}^{M} D_{\mathrm{KL}}(q_m(j) \parallel p(Z_m = j \mid X_i, \phi)) \, , \end{aligned} \tag{S.2}$$

where $D_{\mathrm{KL}}(P \parallel Q)$ is the KL divergence of $P$ and $Q$, so that

$$q_m(j) = p(Z_m = j \mid X_i, \phi) \tag{S.3}$$

implies

$$L(\phi) = \sum_{m=1}^{M} \sum_{j \in J} q_m(j) \log \frac{p(X_m, Z_m = j \mid \phi)}{q_m(j)} \, , \tag{S.4}$$

since $P = Q \Rightarrow D_{\mathrm{KL}}(P \parallel Q) = 0$. Therefore, from an arbitrary parameter guess $\widehat{\phi}^{(t)}$, setting $q_m(j) = p(Z_m = j \mid X_i, \widehat{\phi}^{(t)})$ (the 'E-step') and finding

$$\begin{aligned}
\widehat{\phi}^{(t+1)} &= \arg\max_{\phi} \sum_{m=1}^{M} \sum_{j \in J} q_m(j) \log \frac{p(X_m, Z_m = j \mid \phi)}{q_m(j)} \\
&= \arg\max_{\phi} \sum_{m=1}^{M} \sum_{j \in J} q_m(j) \log p(X_m, Z_m = j \mid \phi) \,, \qquad \text{(S.5)}
\end{aligned}$$

(the 'M-step') guarantees $L(\widehat{\phi}^{(t+1)}) \geq L(\widehat{\phi}^{(t)})$, since

$$\begin{aligned}
L(\widehat{\phi}^{(t+1)}) &\geq \sum_{m=1}^{M} \sum_{j \in J} q_m(j) \log \frac{p(X_m, Z_m = j \mid \widehat{\phi}^{(t+1)})}{q_m(j)} \\
&\geq \sum_{m=1}^{M} \sum_{j \in J} q_m(j) \log \frac{p(X_m, Z_m = j \mid \widehat{\phi}^{(t)})}{q_m(j)} \\
&= L(\widehat{\phi}^{(t)}) \,. \qquad \text{(S.6)}
\end{aligned}$$

**General EM under special conditions** We now consider the special case when the following conditions hold:

$$p(Z_m = j \mid \phi) = \phi_j \,, \qquad \text{(S.7)}$$

$$\sum_{j \in J} \phi_j = 1 \,, \qquad \text{(S.8)}$$

$$p(X_m \mid Z_m = j, \phi) = p(X_m \mid Z_m = j) \,. \qquad \text{(S.9)}$$

This simplifies the M-step,

$$\begin{aligned}
\hat{\phi}^{(n+1)} &= \arg\max_{\phi} \sum_{m=1}^{M} \sum_{j \in J} q_m(j) \log p(X_m, Z_m = j \mid \phi) \\
&= \arg\max_{\phi} \sum_{m=1}^{M} \sum_{j \in J} q_m(j) \log p(X_m \mid Z_m = j) \phi_j \\
&= \arg\max_{\phi} \sum_{m=1}^{M} \sum_{j \in J} q_m(j) \log \phi_j \,, \qquad \text{(S.10)}
\end{aligned}$$

so that the maximising parameter can be found using constrained optimisation. Using Lagrange multipliers, for example,

$$\frac{\partial}{\partial \phi_j} \sum_{m=1}^{M} \sum_{j \in J} q_m(j) \log \phi_j + \beta \left( \sum_{j \in J} \phi_j - 1 \right) = \frac{\sum_{m=1}^{M} q_m(j)}{\phi_j} + \beta \qquad \text{(S.11)}$$

and

$$\begin{aligned}
0 &= \frac{\sum_{m=1}^{M} q_m(j)}{\phi_j} + \beta \\
\iff \qquad \phi_j &= \frac{\sum_{m=1}^{M} q_m(j)}{-\beta} \\
&= \frac{\sum_{m=1}^{M} q_m(j)}{\sum_{m=1}^{M} \sum_{j' \in J} q_m(j')} \,, \qquad \text{(S.12)}
\end{aligned}$$

where the value of $\beta$ is fixed by the constraint that $\phi$s sum to 1.

Finally, we note that under these special conditions, we can rewrite the E-step using Bayes' theorem,

$$\begin{aligned} q_m(j) &= p(Z_m = j \mid X_m, \widehat{\phi}^{(t)}) \\ &= \frac{p(X_m \mid Z_m = j, \widehat{\phi}^{(t)}) p(Z_m = j \mid \widehat{\phi}^{(t)})}{\sum_{j' \in \mathcal{J}} p(X_m \mid Z_m = j', \widehat{\phi}^{(t)}) p(Z_m = j' \mid \widehat{\phi}^{(t)})} \\ &= \frac{p(X_m \mid Z_m = j) \widehat{\phi}_j^{(t)}}{\sum_{j' \in \mathcal{J}} p(X_m \mid Z_m = j') \widehat{\phi}_{j'}^{(t)}} \,. \end{aligned} \tag{S.13}$$

**Multidimensional SFS EM** We now consider the case when the parameter $\phi$ is the (multidimensional) SFS and the latent $Z_m$ is the derived allele count, and simply note that the conditions in equations (S.7) to (S.9) are fulfilled in this case: the probability $p(Z_m = j \mid \phi) = \phi_j$ and $\sum_{j \in \mathcal{J}} \phi_j = 1$ by definition of the SFS, and $p(X_m \mid Z_m, \phi) = p(X_m \mid Z_m)$ by conditional independence of the sequencing data and the SFS given a derived allele count. Therefore, eq. (S.13) and eq. (S.12) recover the E step and M step from the main text.

### References


1. Dempster, A. P., Laird, N. M. & Rubin, D. B. Maximum Likelihood from Incomplete Data Via the EM Algorithm. *Journal of the Royal Statistical Society: Series B (Methodological)* **39,** 1–22 (1977).

### S2 SFS statistics

The results section of the main text refers in several instances to the computation of various statistics from the SFS. The following subsections outline the basis of these calculations.

**Tajima's $\theta$** Based on a one-dimensional SFS $\phi$ from $N$ diploid individuals, Tajima's estimator $\hat{\theta}_{\text{Tajima}}$ of $\theta$ can be calculated [1]

$$\hat{\theta}_{\text{Tajima}} = \binom{n}{2} \sum_{j=1}^{2N-1} j(n-j)\phi_j \,. \tag{S.14}$$

**Hudson's $fst$** For a two-dimensional SFS $\phi$, we can estimate Hudson's $fst$ by [2]

$$\mathrm{F}_{\text{ST}} = \sum_{j \in \mathcal{J} \setminus \{(0,0),(2N_1,2N_2)\}} \frac{(\tilde{p}_{j_1} - \tilde{p}_{j_2})^2 - \frac{\tilde{p}_{j_1}(1-\tilde{p}_{j_1})}{N_1 - 1} - \frac{\tilde{p}_{j_2}(1-\tilde{p}_{j_2})}{N_2 - 1}}{\tilde{p}_{j_1}(1-\tilde{p}_{j_2}) + \tilde{p}_{j_2}(1-\tilde{p}_{j_1})} \phi_j \,, \tag{S.15}$$

where $\mathcal{J} = \{0, \ldots, 2N_1\} \times \{0, \ldots, 2N_2\}$ as in the main text, so that

$$\tilde{p}_{j_i} = j_i / 2N_i \,, \tag{S.16}$$

corresponds to a possible sample allele frequency in the $i$th population.

**$f_2$-statistics** The $f_2$ statistic [3, 4] is defined for two populations,

$$f_2 = \mathrm{E}[(p_1 - p_2)^2] \,, \tag{S.17}$$

where $p_1$, $p_2$ are allele frequencies in the two populations. For a finite sample of $N_1$, $N_2$ individuals from the populations and the two dimensional $\phi$ based on these samples, we estimate

$$\hat{f}_2 = \sum_{j \in \mathcal{J}} (\tilde{p}_{j_1} - \tilde{p}_{j_2})^2 \phi_j \,, \tag{S.18}$$

with $\mathcal{J}$ and $\tilde{p}_{j_i}$ defined as for the $fst$ estimates above.

### References


1. Korneliussen, T. S., Moltke, I., Albrechtsen, A. & Nielsen, R. Calculation of Tajima's D and Other Neutrality Test Statistics from Low Depth Next-Generation Sequencing Data. *BMC Bioinformatics* **14** (2013).
2. Bhatia, G., Patterson, N., Sankararaman, S. & Price, A. L. Estimating and Interpreting $\mathrm{F}_{\mathrm{ST}}$: The Impact of Rare Variants. *Genome Research* **23,** 1514–1521 (2013).
3. Reich, D., Thangaraj, K., Patterson, N., Price, A. L. & Singh, L. Reconstructing Indian Population History. *Nature* **461,** 489–494 (2009).
4. Peter, B. M. Admixture, Population Structure, and F-Statistics. *Genetics* **202,** 1485–1501 (2016).

#### S3 Figures

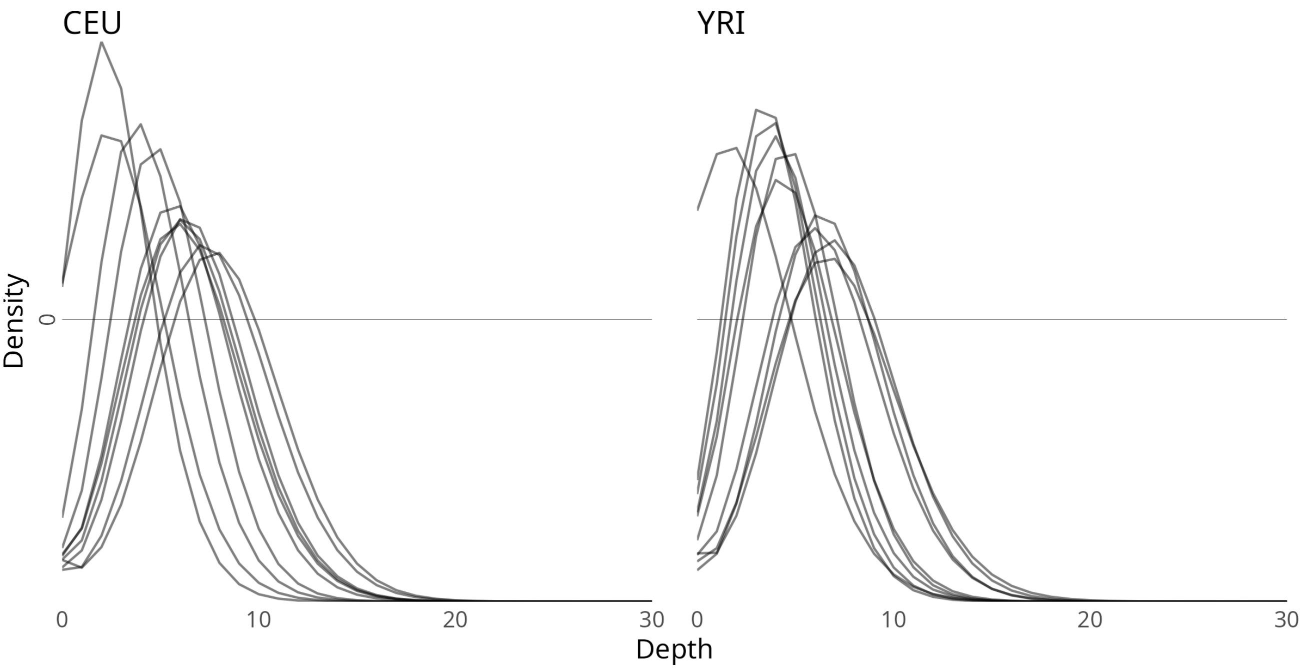


**Supplementary Figure 1:** Human training data depth distributions. Depth over sites for each individual is shown. Sites with depth greater than 30 not shown.

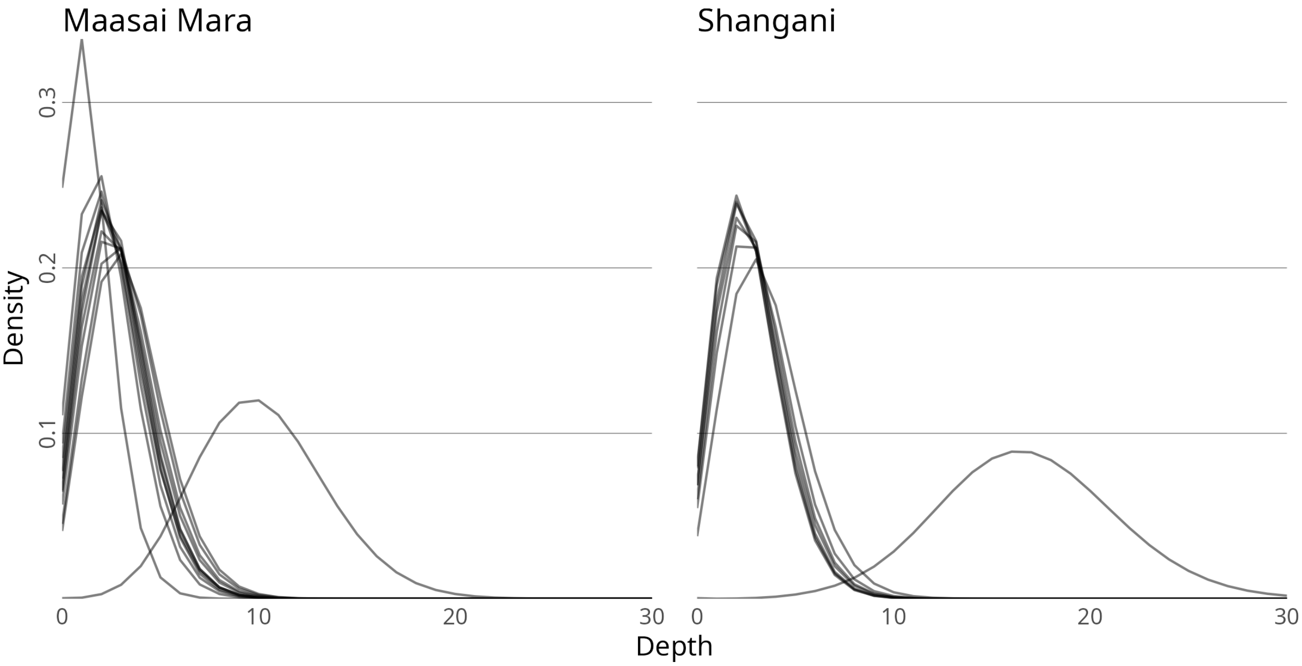


**Supplementary Figure 2:** Impala training data depth distributions. Depth over sites for each individual is shown. Sites with depth greater than 30 not shown.

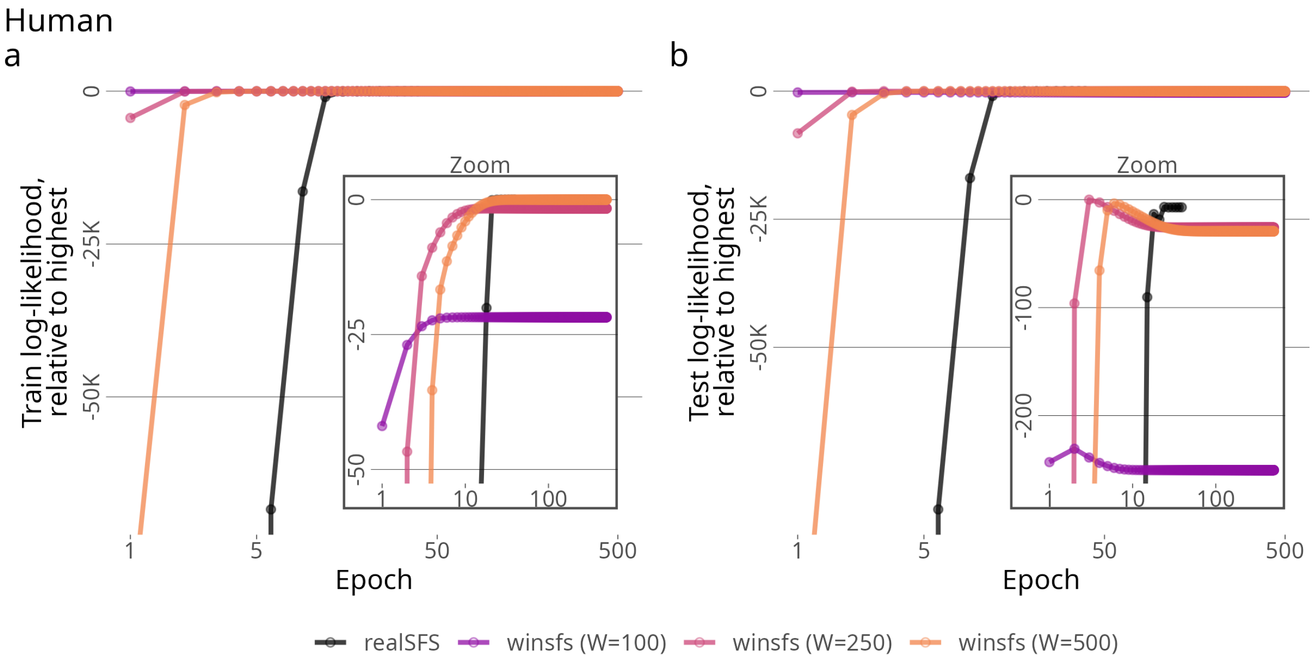


**Supplementary Figure 3:** Log-likelihood over epochs for the human YRI population. **(a)**: Train log-likelihood. **(b)**: Test log-likelihood.

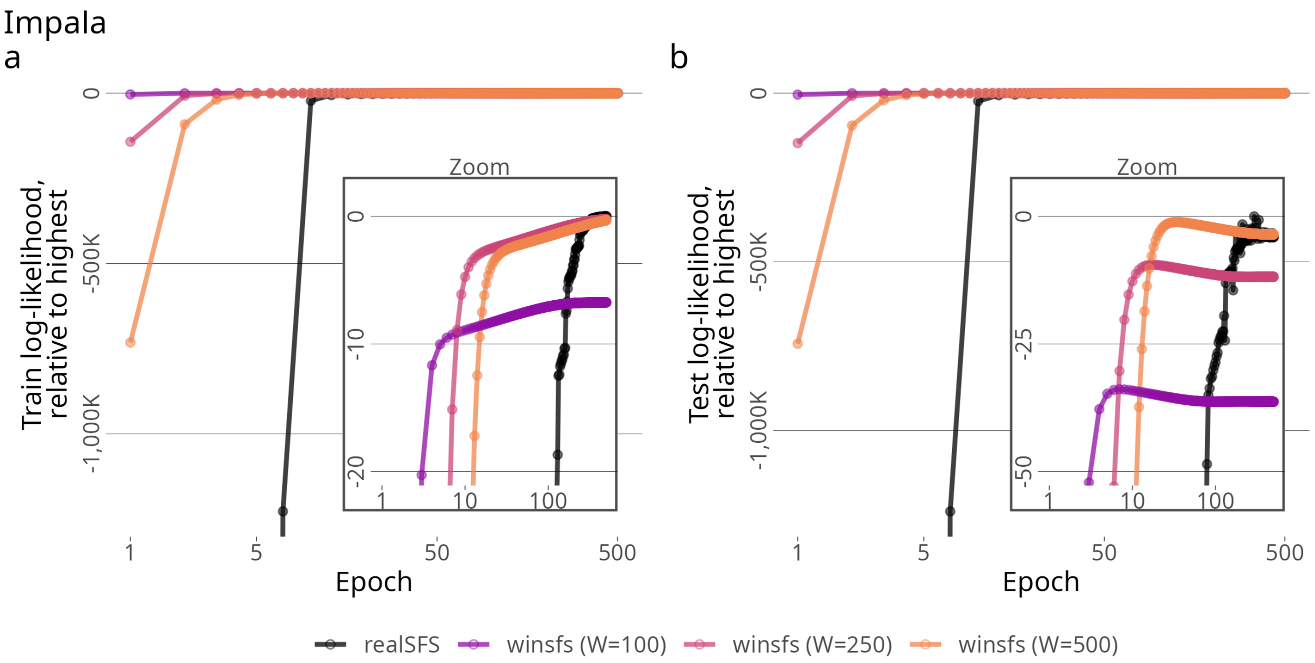


**Supplementary Figure 4:** Log-likelihood over epochs for the impala Maasai Mara population. **(a)**: Train log-likelihood. **(b)**: Test log-likelihood.

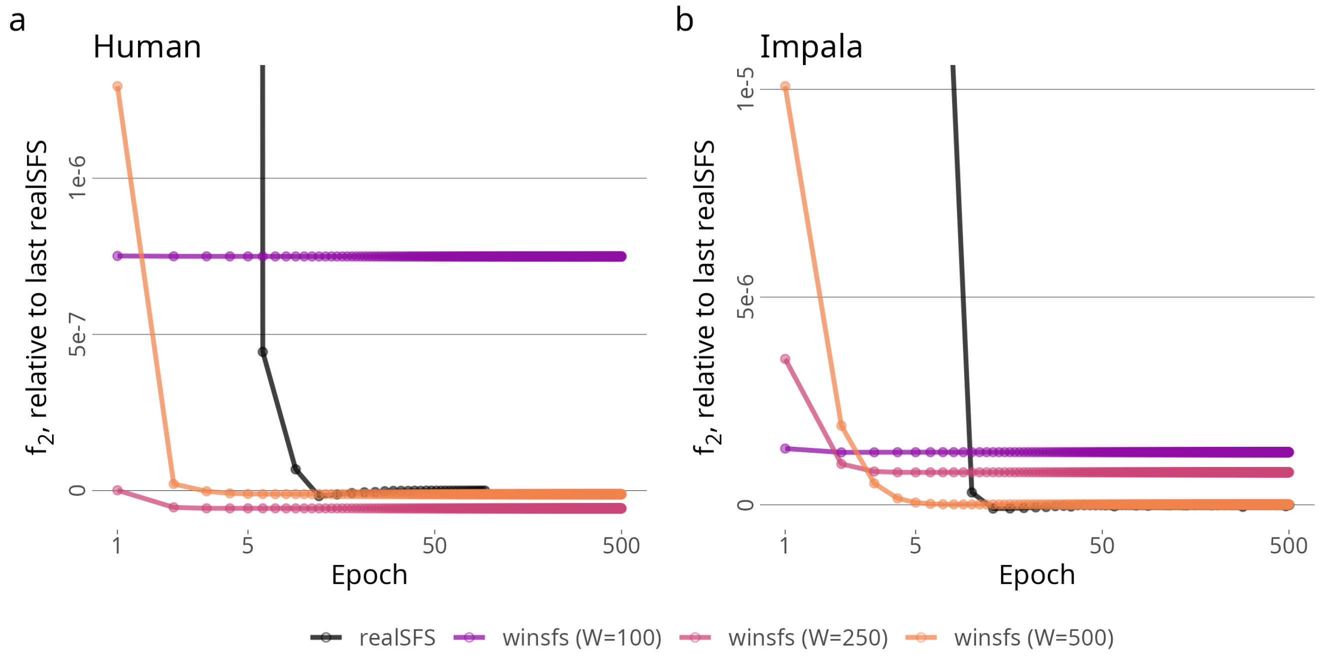


**Supplementary Figure 5:** $f_2$-statistics over epochs. **(a)**: Human CEU/YRI populations. **(b)**: Impala Shangani/Maasai Mara populations.

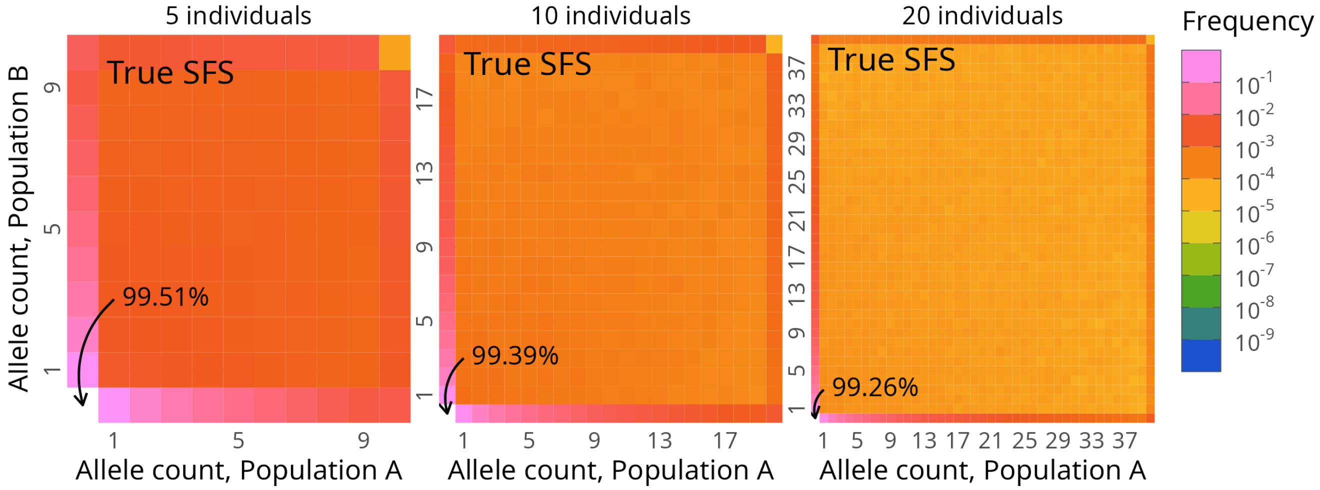


**Supplementary Figure 6:** True spectra for the data simulations. Results shown for scenarios using sample sizes of 5, 10, or 20 individuals. Fixed sites not shown for scale, total proportion indicated by arrows. The colour scale matches the one used in supplementary figure 7 and supplementary figure 8.

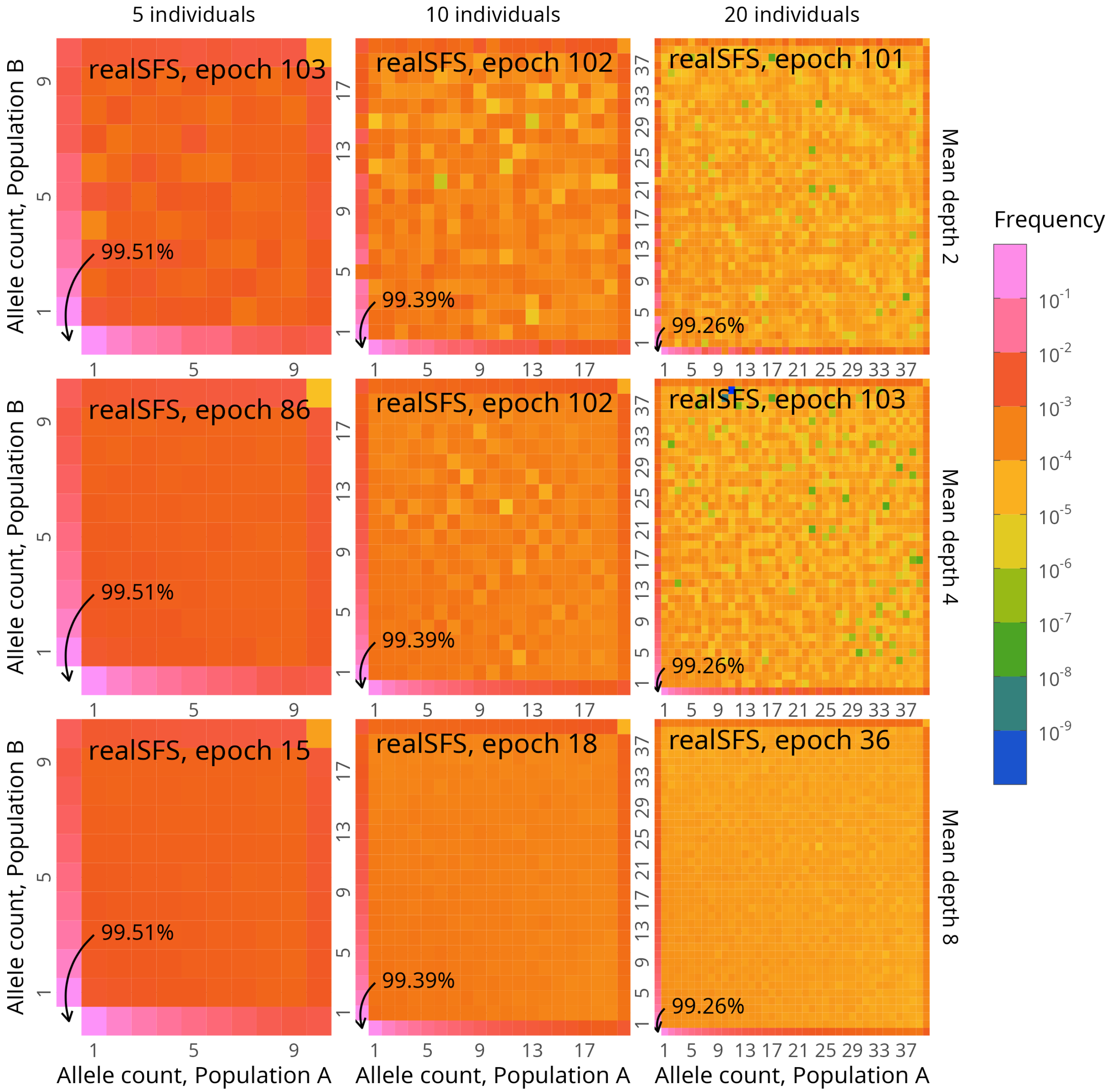


**Supplementary Figure 7:** Spectra inferred by `realSFS` shown at the default stopping time for the simulated data. Results shown for a grid of scenarios using sample sizes of 5, 10, or 20 individuals (labelled top) and mean depth 2, 4, and 8 (labelled right). Panel labels give the stopping epoch. Fixed sites not shown for scale, total proportion indicated by arrows. The colour scale matches the one used in supplementary figure 6 and supplementary figure 8.

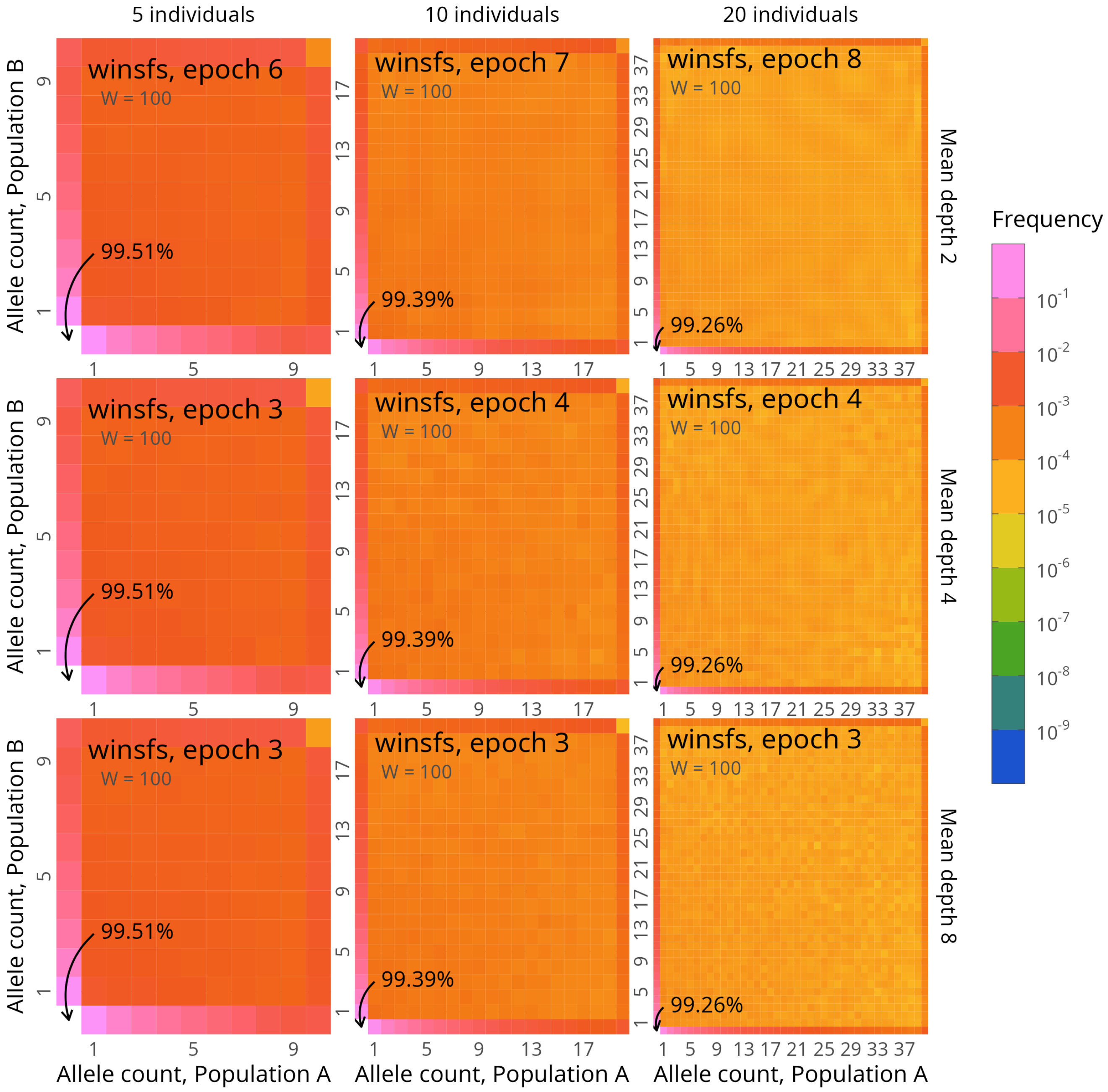


**Supplementary Figure 8:** Spectra inferred by $\texttt{winsfs}_{100}$ shown at the default stopping time for the simulated data. Results shown for a grid of scenarios using sample sizes of 5, 10, or 20 individuals (labelled top) and mean depth 2, 4, and 8 (labelled right). Panel labels give the stopping epoch. Fixed sites not shown for scale, total proportion indicated by arrows. The colour scale matches the one used in supplementary figure 6 and supplementary figure 7.

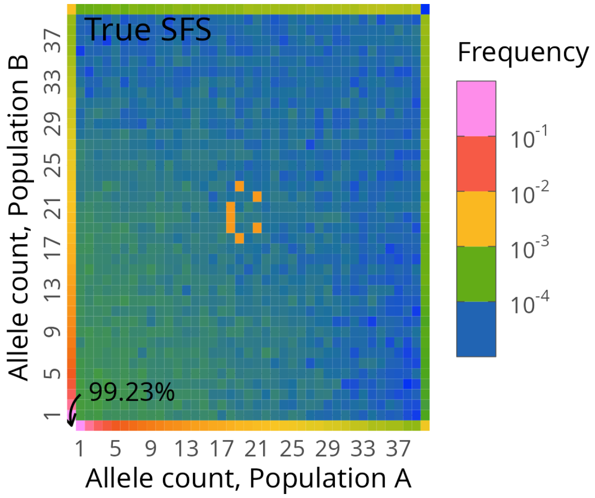


**Supplementary Figure 9:** True spectra for the data simulations with added peaks. This corresponds to the right-most panel of supplementary figure 6, except with 10 000 counts added in seven arbitrary bins near the centre, and without constraining the colour scale. Fixed sites not shown for scale, total proportion indicated by arrows.

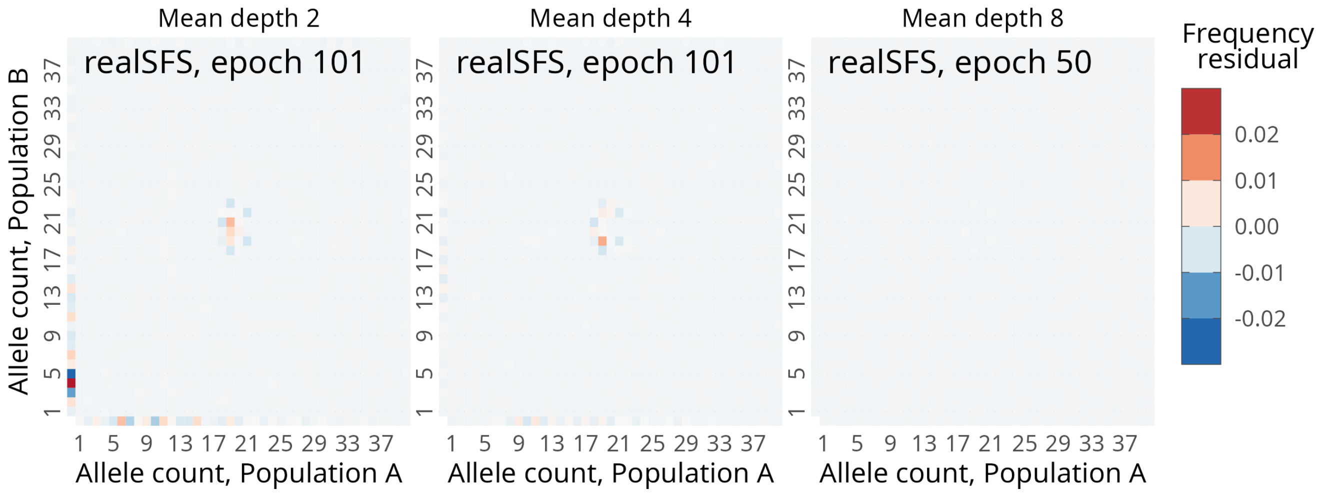


**Supplementary Figure 10:** Residuals of the spectra inferred by `realSFS` shown at the default stopping time for the simulated data with added peaks. The residual is the estimate minus the truth, so that positive values correspond to estimated values being higher than the truth, and negative values to the estimate being lower than the truth. Results shown for a grid of scenarios mean depth 2, 4, and 8. Panel labels give the stopping epoch. The colour scale matches the one used in supplementary figure 11.

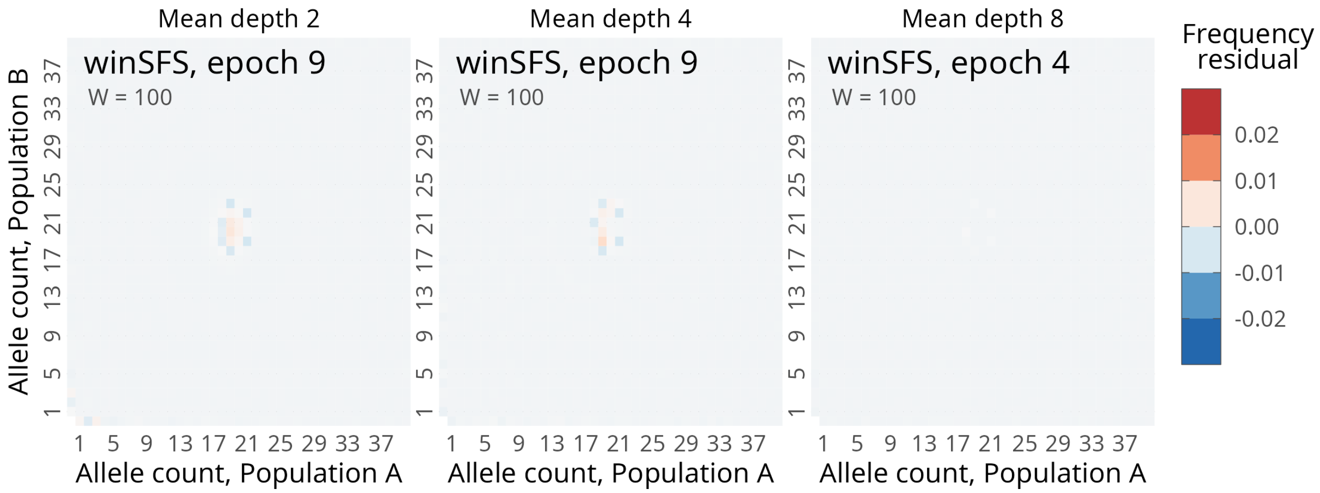


**Supplementary Figure 11:** Residuals of the spectra inferred by `winsfs`$_{100}$ shown at the default stopping time for the simulated data with added peaks. The residual is the estimate minus the truth, so that positive values correspond to estimated values being higher than the truth, and negative values to the estimate being lower than the truth. Results shown for a grid of scenarios mean depth 2, 4, and 8. Panel labels give the stopping epoch. The colour scale matches the one used in supplementary figure 10.

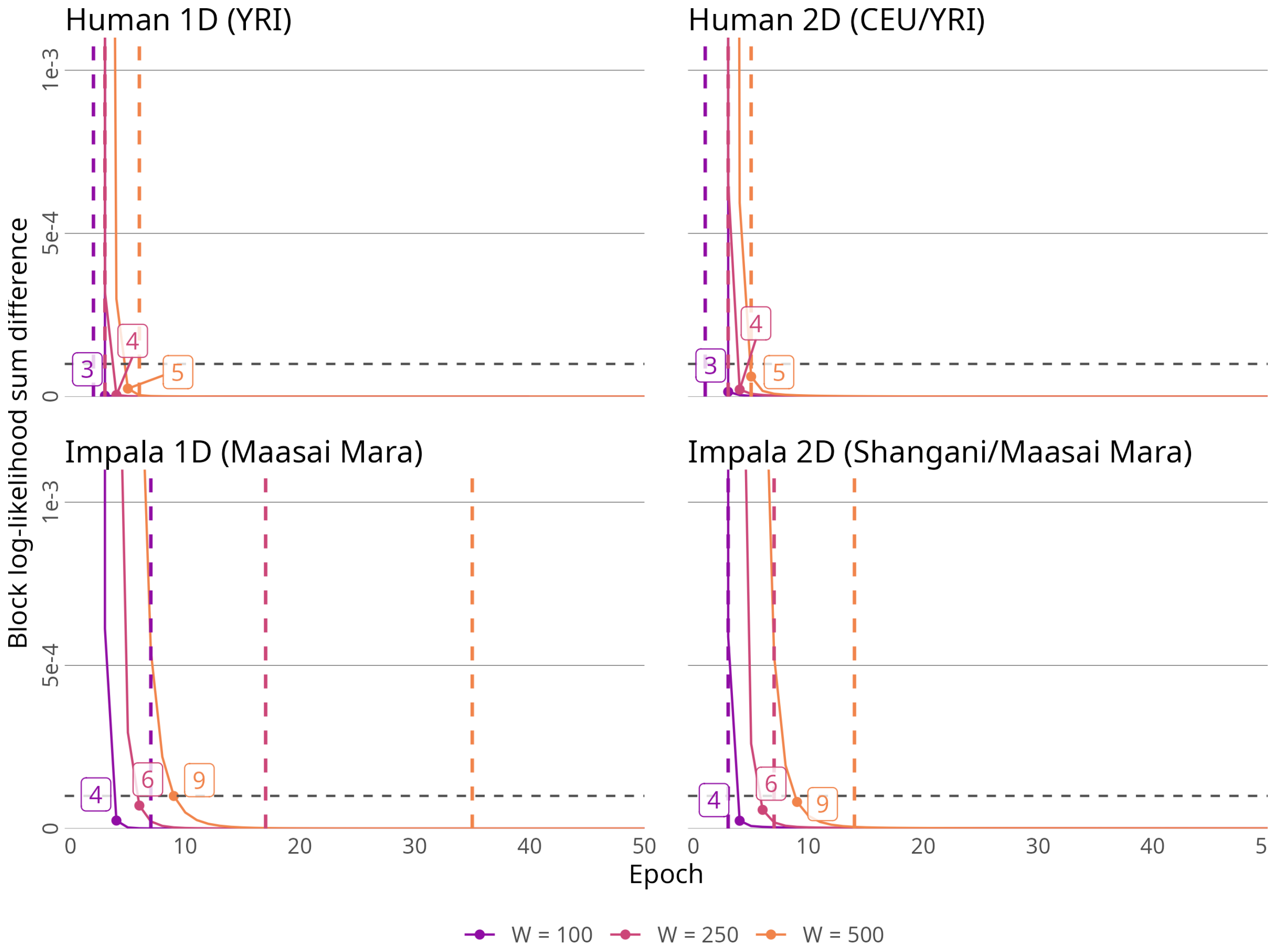


**Supplementary Figure 12:** Difference $L'_e - L_{e-1}$ over epochs $e$ for the different data sets and different window sizes $W \in \{100, 250, 500\}$. The horizontal line marks the chosen default tolerance $\delta = 10^{-4}$, so that the first epoch under the line marks convergence. Number of epochs before convergence is labelled. For convenience, the vertical dashed lines mark the the epoch at which the highest test log-likelihood is achieved.

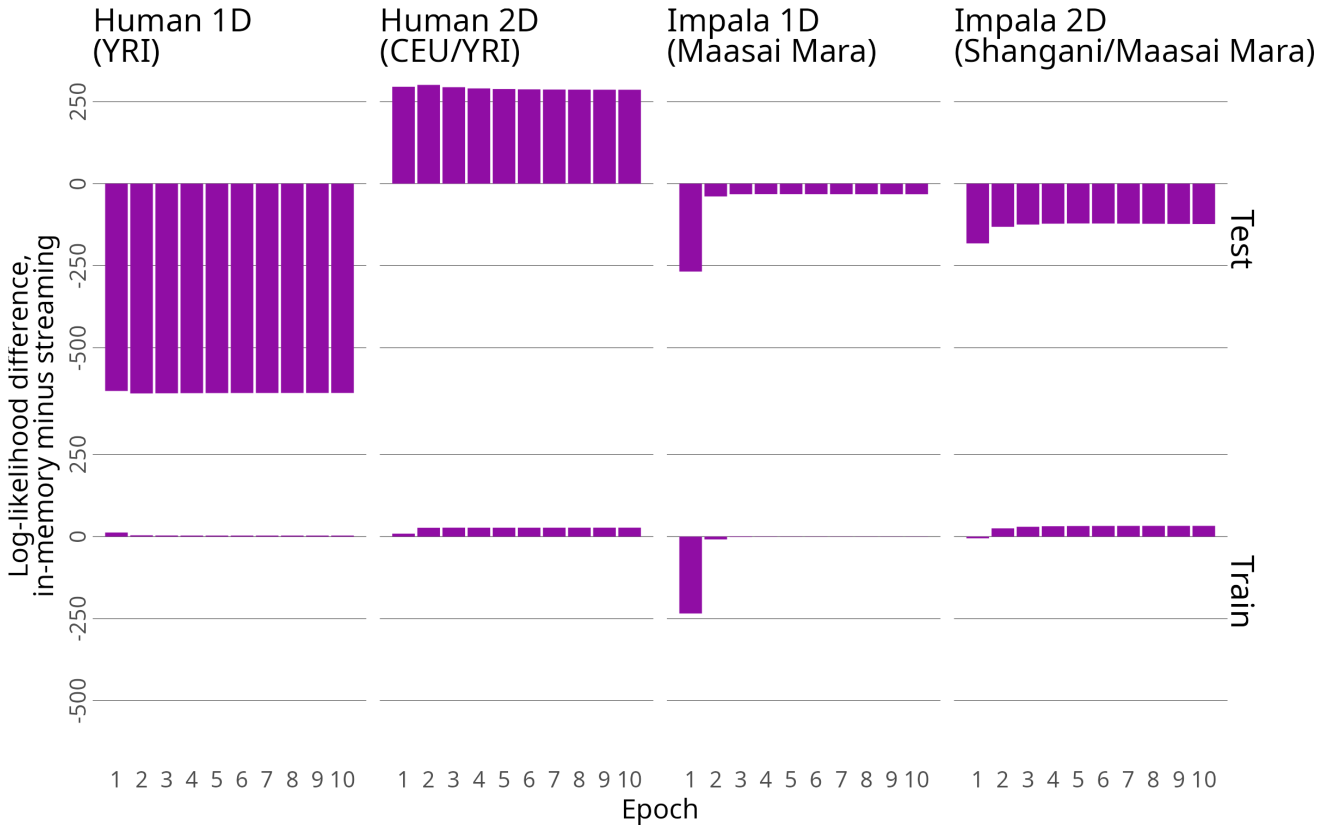


**Supplementary Figure 13:** Comparison of `winsfs` reading the data into memory and streaming pre-shuffled data on the joint impala data set. The difference in train log-likelihood between the two modes is shown for the first five epochs. A positive values means that streaming achieved a higher log-likelihood.

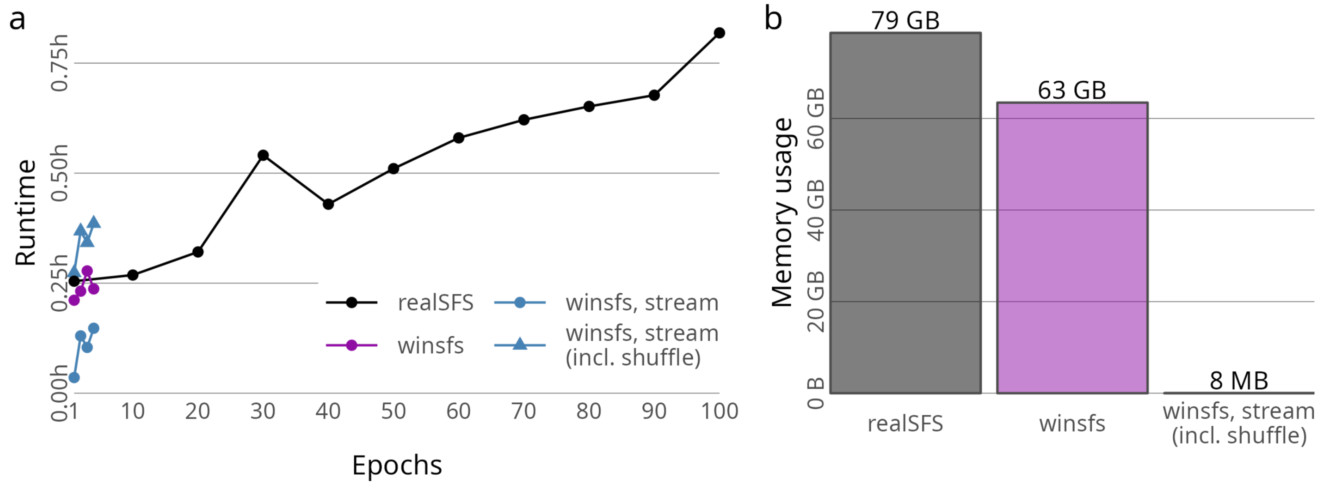


**Supplementary Figure 14:** Computational resource usage of `winsfs` and `realSFS` for the one-dimensional estimation of the Maasai Mara impala population. `winsfs` can be run while loading input data into RAM, or streaming through it on disk. In the latter case, data must be shuffled on disk before hand. **(a)**: Runtime required with 20 threads for various numbers of epochs. Results for `winsfs` are shown for in-memory usage and streaming mode. For streaming modes, times are given with and without the extra time taken to shuffle data on disk before running. **(b)**: Peak memory usage (maximum resident set size).
